# MERTK Coordinates Efferocytosis by Regulating Integrin Localization and Activation

**DOI:** 10.1101/2023.07.20.549870

**Authors:** Brandon H Dickson, Tarannum Tasnim, Austin L Lam, Angela Vreize, Eoin N Blythe, Gregory A Dekaban, Bryan Heit

## Abstract

Efferocytosis – the phagocytic removal of apoptotic cells – is a central component of tissue homeostasis, and in many tissues is mediated by the efferocytic receptor MERTK expressed by macrophages. Although MERTK is critical for efferocytosis, the mechanism by which it directs the engulfment of apoptotic cells is largely unknown. Using immunoprecipitation, mass spectrometry, and super-resolution microscopy, we have identified a pre-formed receptor complex on the macrophage plasma membrane comprised of ∼180 nm clusters of MERTK, β_2_ integrins, and multiple signaling molecules including Src-family kinases, PI3-kinases, and the integrin regulatory proteins ILK and FAK. MERTK is unable to mediate efferocytosis in the absence of β_2_ integrins or their opsonins, while β_2_ integrins require activation via MERTK signaling to induce the engulfment of apoptotic cells. Using FRET microscopy, we determined that MERTK directly induces the conformational change of β_2_ integrins from the low to high-affinity form via a PI3-kinase-dependent signaling pathway. MERTK and β_2_ integrins then form a highly structured synapse in which MERTK is retained by ligand-induced clustering in the synapse centre, while β_2_ integrins and actin form a Src family kinase-, ILK- and FAK-dependent expanding ring which defines the leading edge of the synapse that ultimately engulfs the apoptotic cell. The identification of the MERTK membrane-proximal signaling pathway and the role of β_2_ integrins in this pathway provides new insights into the function of this critical homeostatic receptor and provides new insights into how MERTK mutations and signaling defects may contribute to inflammatory and autoimmune diseases.

## Introduction

The great diversity of phagocytic receptors is a key component of the immune system’s ability to clear pathogens, as this diversity allows phagocytes to engage a broad range of pathogens based on unique pathogen surface chemistry (e.g. dectin-1 binding of fungal glucans), via cooperation with other pathogen-detection systems (e.g. complement-mediated phagocytosis), and to utilize the vast repertoire of the adaptive immune system (e.g. Fc-receptor mediated phagocytosis)[1–3]. Like phagocytosis, efferocytosis – the phagocytic removal of apoptotic cells – is driven by an expansive repertoire of receptors. But unlike phagocytosis, the purpose of this diversity is unclear. Indeed, with only a few exceptions, efferocytic receptors recognize the same ligand – phosphatidylserine (PtdSer, reviewed in [4]). This lipid “eat me” signal is normally restricted to the inner leaflet of the plasma membrane, but during apoptosis, active caspases cleave and activate the membrane scramblase XKR8 while inactivating the flippases ATP11A and ATP11C, thus exposing PtdSer on the outer leaflet [5,6]. While some of this receptor diversity may be explained by cell-type specific expression of some receptors, most efferocytes express multiple efferocytic receptors [4]. For example, macrophages express MERTK, Axl, Tyro3, TIM-4, α_x_ integrin, β_5_ integrin, BAI-1, LRP-1, and CD36 – all receptors reported to bind to apoptotic cells either directly or via opsonins [7–12]. Why a single cell type would need to express such a large repertoire of receptors that recognize the same ligand, especially given that apoptotic cells expose abundant quantities of PtdSer – upwards of 10% of total outer leaflet lipid – remains unclear [13]. One possible explanation for this diversity is the need to closely regulate “multi-purpose” receptors such as integrins. Indeed, both MERTK and TIM-4 have been reported to require integrins for efficient apoptotic cell uptake, relying on the opsonins MFG-E8 and sCD93 to bridge integrins to the apoptotic cell [14–16]. These same integrins, when provided different opsonins or ligands, can also serve as phagocytic receptors, mediators of intravascular adhesion, and as adhesion receptors for cell migration. Some integrins even serve all four roles – for example, α_x_β_2_ mediates efferocytosis via the opsonin soluble CD93 (sCD93), pathogen phagocytosis via the opsonin iC3b, intravascular adhesion via binding to ICAM-2 and VCAM-1, and mediates cell migration [16–19]. The use of integrins for this broad array of activities likely evolved to take advantage of features not found in other adhesion receptors: the ability to regulate their affinity via conformational changes, the high affinity of integrins for their ligands, and the ability of integrins to engage actin-based force-generating machinery within the cell [20–22].

This diverse range of activities requires that integrins be carefully regulated. This regulation generally occurs via other receptors whose signaling regulates integrin affinity through inside-out signaling, allowing these receptors to direct the resulting pattern of integrin activation. Antibody-mediated phagocytosis via Fcγ receptors (FcγR) shares many mechanical similarities to efferocytosis and may provide insights into how integrins are regulated during efferocytosis. During the phagocytosis of IgG-opsonized targets, FcγR signaling induces the formation of a phagocytic synapse in which FcγRs are concentrated in the centre and are surrounded by a bounding ring of active α_m_β_2_ integrin (CD11b/CD18, Mac-1) [23]. This outer ring of α_m_β_2_ does not appear to serve an adhesive or force-generating role, and instead forms a diffusion barrier that excludes the phosphatase CD45. This exclusion of CD45 from the phagocytic cup ensures uninterrupted Src-family kinase (SFK) signaling, which regulates the contractile forces required for engulfment of the antibody-opsonized pathogen [24]. In contrast, during complement-mediated phagocytosis, α_m_β_2_ and α_x_β_2_ bind to complement iC3b, which is deposited on the surface of pathogens following activation of the complement cascade [2,25]. Unlike in FcγR-mediated efferocytosis, the integrins are the primary phagocytic receptor in this form of phagocytosis, providing both adhesion to the target, and generating the actin-mediated contractile force that draws the pathogen into the phagocyte [26,27]. Clearly, the roles of integrins in phagocytosis-like processes can vary depending on the ligands present on the phagocytic target, and on the signaling being generated by other receptors within or near the phagocytic synapse.

MERTK is the sole or predominant efferocytic receptor in multiple tissues including the eye, brain and heart, and is highly expressed in macrophages – cells which function as the primary efferocyte in many tissues [8,15,28]. Single nucleotide polymorphisms in MERTK and its opsonins are associated with a range of diseases including retinitis pigmentosa, male infertility, autoimmune disorders such as multiple sclerosis and systemic lupus erythematosus, neurological disorders including Alzheimer’s disease, and inflammatory diseases including atherosclerosis [29–32]. MERTK is bridged to PtdSer on apoptotic cells by two well-characterized opsonins: Gas6 and Protein S. Engagement of an apoptotic cell activates MERTK’s intrinsic tyrosine kinase domain, initiating a signaling pathway whose membrane-proximal signaling is incompletely understood, but which converges on the adaptor Grb2, phosphatidylinositol-3-kinase (PI3K), SFKs, and Vav3 [33]. In retinal pigment epithelial cells, this pathway engages α_v_β_5_ integrin to mediate the efferocytosis of shed photoreceptor outer segments, with the α_v_β_5_ proposed to activate Rac1, which in turn mediates the actin reorganization necessary to engulf the apoptotic cell [34–36]. However, the role of integrins in MERTK-mediated efferocytosis is controversial, with some studies suggesting that integrin binding is a pre-requisite for MERTK activation, others suggesting that MERTK activates integrins, and yet other studies suggesting that integrins are dispensable for MERTK-mediated phagocytosis [15,37–39].

Using mass spectrometry, selective engagement of MERTK and β_2_ integrins, and quantitative microscopy, we demonstrate that MERTK signaling coordinates the formation a highly ordered synapse-like structure between macrophages and apoptotic cells, with MERTK regulating synapse structure and the positioning and activation of β_2_ integrins within the synapse. This coordinated signalling produces an expanding integrin ring which engulfs the apoptotic cell, with efficient macrophage efferocytosis requiring this coordinated activity of MERTK and β_2_ integrins.

## Results

### Macrophage Efferocytic Receptor Expression and Usage

PMA-differentiated THP-1 macrophages expressed a broad range of efferocytic receptors, including all three members of the TAM family (**Figure 1A**), receptors which directly bind to PtdSer (**Figure 1B**), integrins (**Figure 1C**), scavenger receptors (**Figure 1D**), and inhibitory receptors (**Figure 1E**). Apoptotic Jurkat cells and apoptotic mimics comprised of 3 µm beads coated in a 20:80 mixture of PtdSer:phosphatidylcholine (PtdChol) were opsonized with serum – which contains apoptotic cell opsonins for integrins (MFG-E8 and complement) and TAM receptors (Gas6 and ProS), and were used to determine the relative role of the TAM receptors in macrophage efferocytosis. Despite the wide array of efferocytic receptors expressed in these cells, siRNA knockdown of MERTK abrogated efferocytosis of both the mimics and apoptotic cells, with even a triple-TAM knockdown having no additive effect over MERTK-knockdown alone **(Figures 1F-G, S1A**). A similar trend was observed when THP-1 macrophages were treated with the MERTK kinase domain inhibitor, UNC2250 **(Figure 1H**) [40]. While these results do not eliminate potential roles for the other receptors at later stages of apoptotic cell engulfment, they indicate that at a minimum, macrophages rely on MERTK for the initial recognition of apoptotic cells. Given this central role of MERTK in the initial recognition of apoptotic cells, we further probed MERTK organization on macrophages.

**Figure 1:**
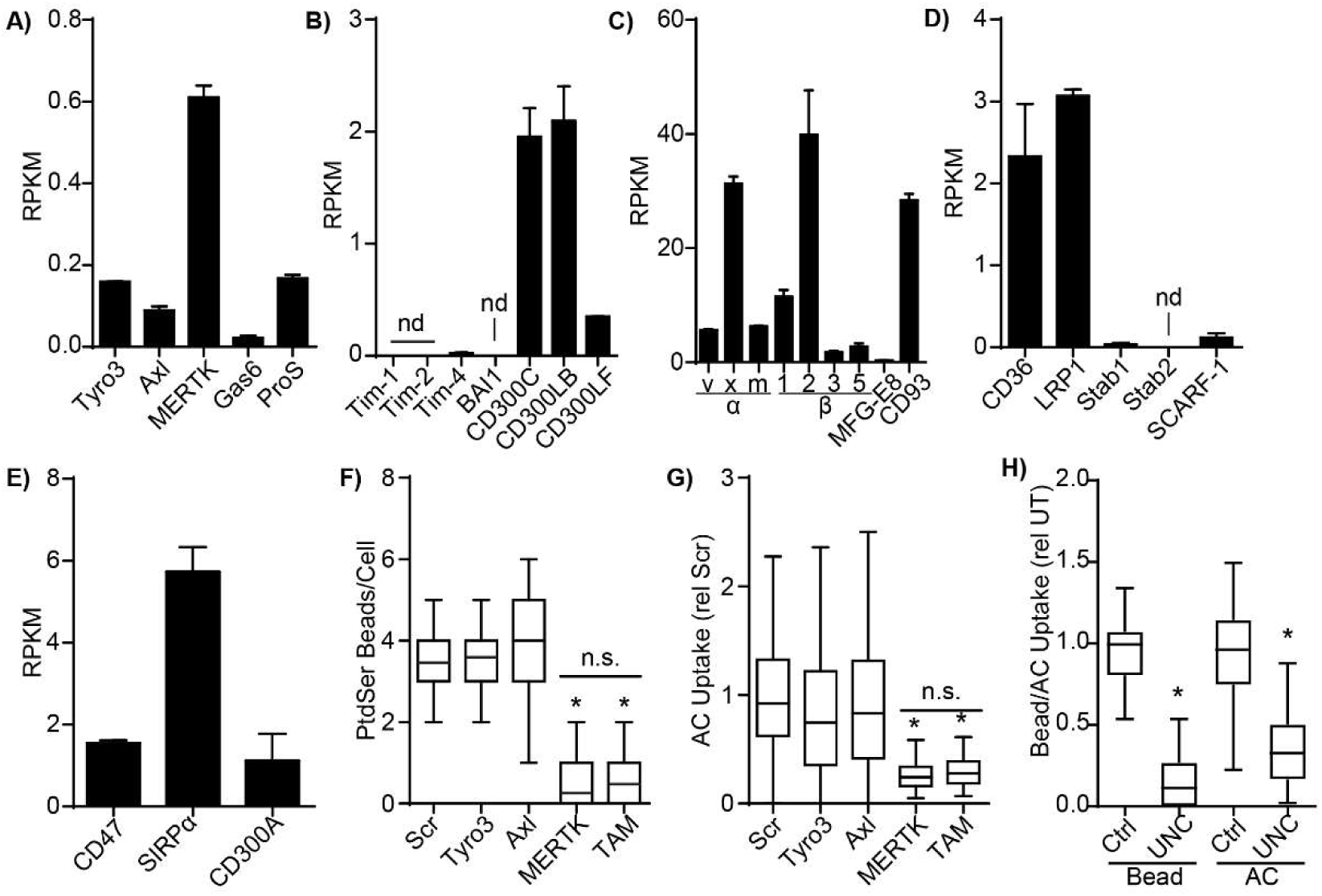
Efferocytic Receptor Expression and Function in THP-1 Macrophages. **A-E)** mRNA expression of (A) TAM-family efferocytic receptors (Tyro3, Axl and MERTK) and their opsonins Gas6 and Protein S (ProS), (B) PtdSer-binding efferocytic receptors, (C) α-chains (α_v_, α_x_, α_m_), β-chains (β_1_, β_2_, β_3_, β_5_), and the opsonins (MFG-E8 and CD93) of efferocytic integrins, (D) scavenger receptors known to bind to apoptotic cells, and (E) inhibitory efferocytic receptors were quantified in RNAseq data from PMA-differentiated THP-1 macrophages. n.d. = not detected. **F-G)** Quantification of the efferocytosis of serum-opsonized apoptotic mimics (F) or apoptotic cells (G) by THP-1 macrophages where Tyro3, Axl, MERTK or all three receptors (TAM) were siRNA-depleted. Scr = scrambled siRNA control. **H)** Efferocytic uptake of apoptotic mimics and apoptotic Jurkat T cells by macrophages treated with 5 µM of the MERTK inhibitor UNC2250 (UNC) or vehicle control (Ctrl). Data is plotted as mean ± SEM (A-E) or median ± interquartile range with whiskers indicating the 95% confidence interval (F-H). n = 3, * = p < 0.05, n.s. = p > 0.05 compared to Scr (F-G) or UT (H), Kruskal-Wallis test with Dunn’s multiple comparisons test.

### Characterization of the MERTK Signalosomes

We previously determined that human MERTK is highly clustered within regions ∼180 nm in diameter on the surface of resting human macrophages [41], suggesting that MERTK exists in pre-formed signalosomes prior to efferocytosis. To determine if other proteins were co-clustered in these MERTK signalosomes, we expressed MERTK with an extracellular HA-tag, then used reversible cross-linking to immunoprecipitate MERTK and any attendant proteins. Immunoprecipitation/mass spectrometry identified several proteins associated with MERTK on resting cells, including β_1_ and β_2_ integrins, integrin-associated signalling molecules (FAK, ILK), members of the PI3K signalling pathway, regulators of Rho-family GTPases, and several kinases including multiple SFKs (**Figure 2A**, **Table 1**). To validate these interactions and investigate the spatial organization of the immunoprecipitated proteins with MERTK, spatial statistics were applied to images with ∼20 nm resolution gathered using Ground State Depletion Microscopy (GSDM). As is common with plasma membrane proteins, most of the detected proteins exhibited significant self-clustering (**Figure 2B, S2**), with the bulk of MERTK clustered into structures ∼180 nm in diameter, similar to what we reported earlier. Radial distribution analysis identified an unexpectedly complex spatial relationship between MERTK and the β_1_, β_2_, and α_x_ integrins (**Figure 2C**). β_1_ integrin was strongly enriched at distances from 20 to 100 nm from MERTK, indicating that MERTK and β_1_ integrin are co-clustered in regions 100-200 nm in diameter. While a portion of β_2_ and α_x_ integrins co-clustered with MERTK in regions of similar diameter, a secondary enrichment of β_2_ and α_x_ integrins was found at radii of 100 nm to 300 nm—indicating that in addition to being co-clustered with MERTK, that clusters of these integrins also contact MERTK clusters without intermixing (**Figure 2C, S2**).

**Figure 2:**
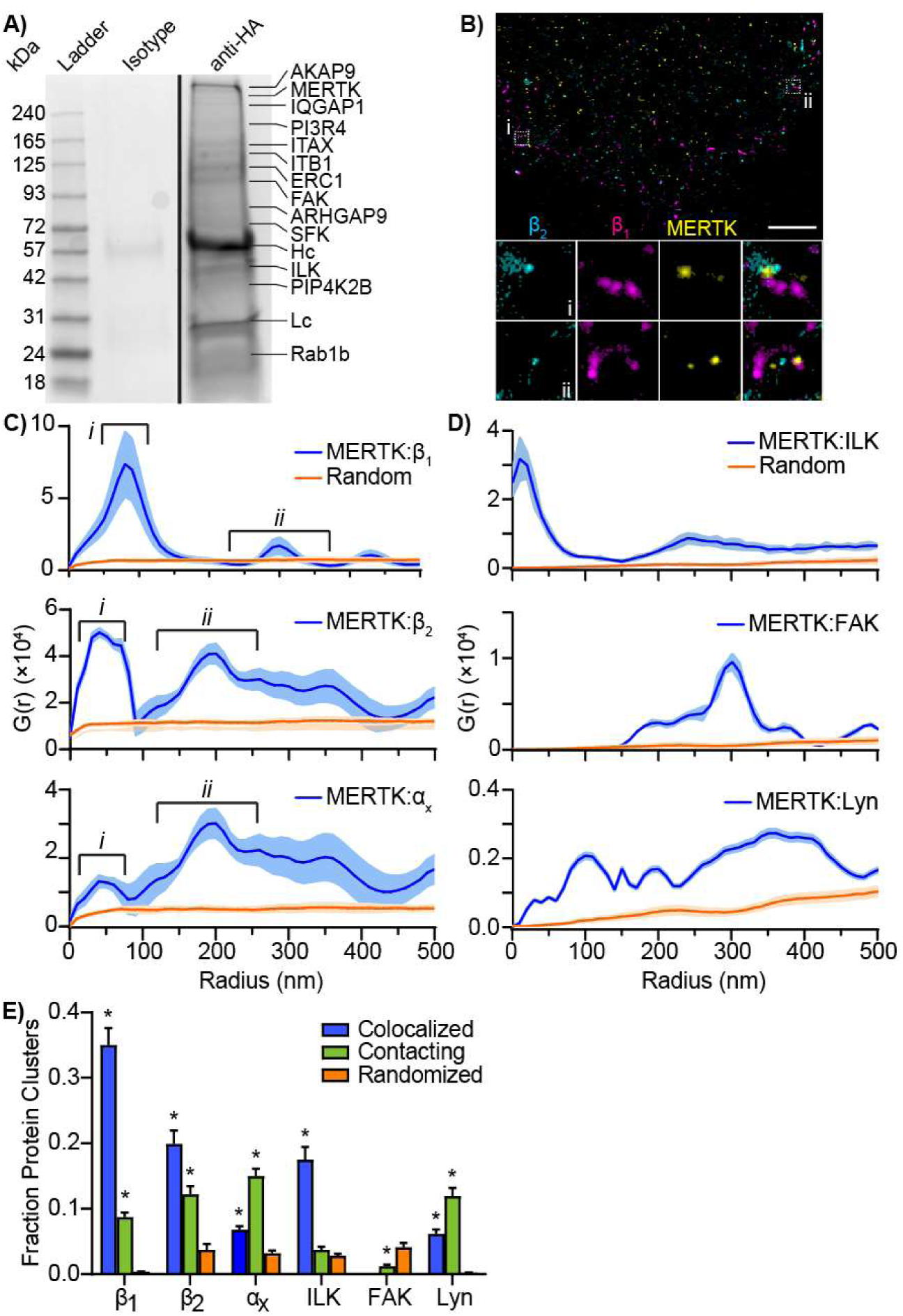
Identification of the MERTK Signalosome. **A)** Representative anti-HA-MERTK immunoprecipitation. Reversible crosslinking and mass spectrometry were used to identify proteins which co-immunoprecipitate with MERTK. Isotype = irrelevant antigen antibody, anti-HA = HA-MERTK immunoprecipitation. Major proteins found in each band are identified, Hc = heavy chain of precipitating antibody, Lc = light chain of precipitating antibody. **B)** Representative GSDM image of MERTK (Yellow) and β_1_ integrin (Magenta) and β_2_ integrin (Cyan) on the macrophage surface. Insets are of two dashed box (i and ii) showing MERTK co-clustered with both β_1_ and β_2_ (i) and co-clustered with only β_1_ (ii). Scale bar is 2 µm. **C)** Radial distribution analysis of the density of β_1_, β_2_, and α_x_ integrins as a function of their distance from MERTK (blue). The integrins shows two distinct clusters with MERTK: (i) co-clustering within the ∼180 nm diameter MERTK clusters, and (ii) clusters which touch, but are not intermixed with, the MERTK clusters. Random (orange) = G(r) measured when the position of MERTK has been randomized. **D)** Radial distribution analysis of the co-clustering of ILK, FAK and Lyn with MERTK. Random (orange) = G(r) measured when the position of MERTK has been randomized. **E)** Fraction of protein clusters that are intermixed with MERTK clusters (Colocalized) or contacting but not intermixed with MERTK clusters (Contacting). Randomized = Fraction of clusters which contact or overlap with MERTK when the position of MERTK clusters are randomized. All clusters were identified in GSDM images using the OPTICS algorithm. n = 3-5, * = p < 0.05 compared to Randomized, ANOVA with Tukey correction.

**Table 1:** Proteins Identified by Mass Spectrometry Following MERTK Immunoprecipitation.

| Gene Symbol | Peptides | Coverage (%) | p |
| --- | --- | --- | --- |
| *Integrins* |  |  |  |
| ITB1 ($\beta_1$ integrin) | 15 | 21 | $9.43 \times 10^{-4}$ |
| ITAX ($\alpha_x$ integrin) | 55 | 22 | $5.47 \times 10^{-4}$ |
| ILK | 6 | 8 | $1.19 \times 10^{-3}$ |
| FAK | 3 | 3 | $1.44 \times 10^{-2}$ |
| *GTPases* |  |  |  |
| IQGAP1 | 18 | 31 | $5.49 \times 10^{-4}$ |
| RhoE | 6 | 2 | $2.66 \times 10^{-3}$ |
| ARHGAP9 | 4 | 5 | $2.04 \times 10^{-3}$ |
| *Src-Family Kinases* |  |  |  |
| Lyn | 2 | 2 | $3.51 \times 10^{-2}$ |
| Fgr | 3 | 2 | $2.63 \times 10^{-2}$ |
| Hck | 2 | 3 | $3.42 \times 10^{-2}$ |
| Fyn | 4 | 3 | $1.78 \times 10^{-2}$ |
| *Phosphatidylinositol Signalling* |  |  |  |
| PI3R4 | 4 | 4 | $1.14 \times 10^{-3}$ |
| PIP4K2B | 3 | 1 | $2.99 \times 10^{-2}$ |
| PI42C | 8 | 3 | $1.34 \times 10^{-3}$ |
| *Protein Kinases* |  |  |  |
| Erc1 | 8 | 7 | $1.47 \times 10^{-3}$ |
| MAP4K4 | 5 | 5 | $1.06 \times 10^{-3}$ |
| RASA2 | 2 | 1 | $4.31 \times 10^{-2}$ |
| AKAP9 | 2 | 3 | $1.87 \times 10^{-2}$ |

This analysis was then performed on the signaling molecules ILK, Lyn, and FAK, with the first two proteins co-clustering with MERTK in a pattern reminiscent of the integrins (**Figure 2D, S2**). Unexpectedly, FAK was excluded from MERTK clusters, suggesting that FAK likely co-precipitated with MERTK-associated integrins rather than with MERTK itself. To ensure that these were bona fide interactions, and not merely coincidental interactions creating the illusion of co-clustering/contacting, the positions of all molecules in the image were randomized over the same area and the co-clustering analysis repeated, with this analysis demonstrating that the integrin-MERTK co-clusters and contacting clusters occurred at a frequency 50 to 2,000 times that expected of randomly distributed proteins (**Figure 2C-D**). To further characterize the interaction of MERTK with these proteins, individual clusters of MERTK and these proteins were identified by OPTICS segmentation and the fraction of each protein which overlapped with a cluster of MERTK, or physically contacting but not intermixed with MERTK, determined (**Figure 2E**).

Consistent with the radial distribution analysis, all three integrins interacted extensively with MERTK, as did ILK and Lyn. These interactions were not due to chance positioning of protein clusters, as the observed degree of overlapping and contacting clusters was greatly decreased when the position of MERTK clusters were randomized over an equivalent area (**Figure 2E**). Combined, these data demonstrate that MERTK is structured into preformed 180 nm diameter signalosomes which also contain several integrins, ILK, and the Src-family kinase Lyn.

### MERTK Forms an Efferocytic Synapse via a Novel Signaling Pathway

Given the high degree of MERTK-α_x_ integrin co-clustering, and previous reports indicating that MERTK requires integrins for its function, we next investigated the roles of MERTK and α_x_β_2_ integrin in efferocytosis, using apoptotic cell mimics opsonized either with the MERTK opsonin Gas6, the α_x_ integrin opsonin sCD93, or both opsonins, with both the binding and internalization of the mimics quantified (**Figure 3A**). Consistent with our observation that MERTK is the predominant efferocytic receptor on macrophages (**Figure 1F-G**), neither binding nor internalization occurred when mimics lacked the MERTK opsonin Gas6, or when MERTK was siRNA depleted. Mimics were bound by macrophages when both Gas6 and MERTK were present, but engulfment required the additional presence of both sCD93 and α_x_ integrin (**Figures 3A, S1B**). This suggests that MERTK activates integrins upon recognition of an apoptotic cell, with the integrins then providing the necessary affinity and/or physical forces required for engulfment. Consistent with this model, mimics bearing only the α_x_ integrin ligand sCD93 were bound and internalized when integrins were forced into their high-affinity conformation by the addition of manganese (**Figure 3A**).

**Figure 3:**
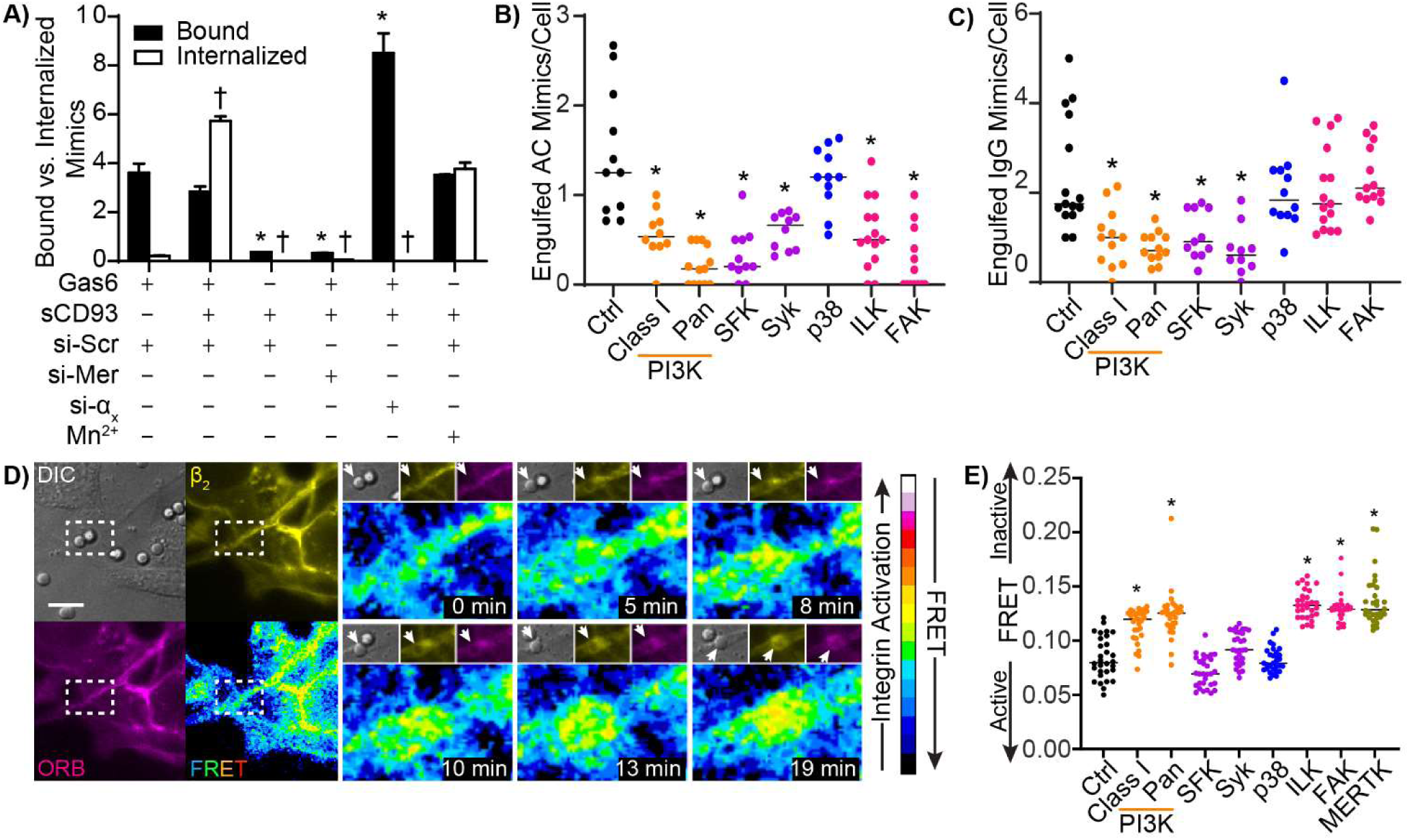
MERTK Regulates Integrin Affinity Through a Novel Signaling Pathway. **A)** Binding and internalization of apoptotic cell mimics opsonized with the MERTK opsonin Gas6, the α_x_ integrin opsonin sCD93, or both opsonins, by THP-1 macrophages that are either WT, treated with a scrambled (non-targeting) siRNA (si-Scr), MERTK-depleting siRNA (si-Mer), α_x_ integrin-depleting siRNA (si-α_x_), or where integrins are forcibly activated by addition of 1 mM MgCl_2_ (Mg^2+^). **B-C)** Engulfment of apoptotic cell mimics (B) and IgG-coated phagocytic targets (C) by THP-1 macrophages treated with inhibitors of Class I PI3K (LY294002), Pan-PI3K (Wortmanin), SFK’s (PP1), Syk (Syk-I), p38 MAPK (SB203580), ILK (Cpd-22), FAK (PF-00562271), or a vehicle control (Ctrl). **D)** Representative FRET time-lapse of β_2_ integrin activation in response to MERTK-specific (Gas6-opsonized) apoptotic cell mimics. *Left:* low-magnification field of the region under observation at 0 min showing white-light (DIC), β_2_ integrin (yellow), ORB (magenta), and FRET channels. *Right:* Time-series of the region indicated by the white box, showing an increase in integrin activation beneath the apoptotic cell mimicking bead (arrows). Scale bar is 10 µm, insert is 14.86 × 10.29 µm. **E)** β_2_ integrin activation in THP-1 macrophages treated with the same inhibitors as in panels B-C, plus an inhibitor of MERTK (UNC2250), as measured by FRET microscopy. n = a minimum of 30 cells/condition imaged in 3 independent experiments. * = p < 0.05 compared to bound Gas6-opsonized beads bound by wild-type macrophages (A) or vehicle control macrophages (B,C,E), † = p < 0.05 compared to Gas6-opsonized beads internalized by wild-type macrophages, Kruskal-Wallis test with Dunn’s multiple comparisons test.

Based on our mass spectrometry results (**Table 1**) and known phagocytic signalling pathways [42], we identified a number of signaling molecules potentially important for the MERTK-mediated efferocytosis—specifically, PI3K, SFKs, Syk (downstream of SFKs), p38 MAPK (downstream of MAP4K4), and the integrin-associated signalling molecules ILK and FAK. Using pharmacological inhibitors of these proteins and serum-opsonized apoptotic cell mimics, we determined that all but p38 MAPK were required for efficient engulfment (**Figure 3B**). In comparison, the phagocytic engulfment of IgG-opsonized pathogen mimics required only PI3K, SFKs, and Syk (**Figure 3C**). Thus, MERTK mediates efferocytosis via a signaling pathway that diverges from the canonical phagocytic FcγR signaling pathway [42]. We next probed the role of these pathways in regulating the activation of β_2_ integrins using sensitized emission Fluorescence Resonance Energy Transfer (FRET) between a β_2_ integrin headgroup-specific FITC-labeled antibody and the membrane dye octadecyl-rhodamine B (ORB) [21]. In the low affinity (bent) conformation, the antibody will be located near the membrane, allowing for FRET to occur between FITC and the ORB in the cell membrane. Activation of the integrin induces a conformational change into an extended high-affinity confirmation, thus moving the headgroup out of the range of FRET, producing a loss of FRET signal (**Figure S3**). Prior to contacting Gas6-opsonized apoptotic cell mimic, extensive FRET is observed in the macrophage membrane, indicating that the β_2_ integrins are largely in an inactive conformation (**Figure 3D**). As the macrophage engages the bead, a phagocyte cup-like structure forms around the apoptotic cell mimic, and a loss of FRET signal (indicating an activation of integrins) is observed (**Figures 3D, S4, Video S1**). As these beads lack any integrin opsonins, this integrin activation must occur via Gas6-MERTK signaling. Interestingly, MERTK-mediated integrin activation within the efferocytic cup required PI3K, ILK, FAK, and the kinase activity of MERTK, but not SFKs or Syk (**Figure 3E**), suggesting that the requirement of SFKs and Syk for efferocytosis are restricted to integrin-independent aspects of efferocytosis.

A “frustrated efferocytosis” model—wherein THP-1 macrophages are “parachuted” onto planar supported lipid bilayers comprised of the same lipid mixture as our apoptotic cell mimics—was used to investigate the spatial organization of MERTK and β_2_ integrin during efferocytosis [43,44]. Macrophages form an efferocytic synapse when they contact these planar apoptotic cell mimics (**Figure 4A, Video S2**), with formation of this synapse dependent on the presence of PtdSer and appropriate opsonins, dependent on all tested signaling molecules, and was dependent on integrin activation (**Figure S5A-C**). Initially, MERTK and β_2_ integrin are colocalized, but very quickly move into separate spatial domains, with the integrins separating from MERTK and forming an expanding ring at the edge of the synapse within 6 minutes of synapse formation **(Figure 4B**). Interestingly, this appears to occur via two mechanisms. Firstly, MERTK and β_2_ integrins from the adjacent membrane are incorporated into the synapse as the synapse expands. Once in the synapse, MERTK remains close to its original location within the synapse, while β_2_ integrins are retained at the expanding outer edge (**Figure 4A, Videos S2 & S3**). Secondly, at later timepoints β_2_ integrins – but not MERTK – are exocytosed within the synapse itself (arrows in Figure 4A & 4D). As we only label extracellular integrins at the start of the assay, these integrins must have been endocytosed elsewhere on the macrophage, trafficked to the synapse, and then exocytosed. The expansion of the synapse is dependent on PI3K, SFKs, FAK, ILK and an active MERTK kinase domain (**Figure 4C**), but interestingly did not require Syk. This unexpected, as in canonical FcγR-mediated phagocytosis Syk is central to the induction of the actin reorganization required to engulf bound pathogens [45]. We quantified β_2_ integrin activation within the synapse using FRET. At early timepoints integrins are in their inactive state, as were recently exocytosed integrins, but integrins rapidly activated as the synapse forms (**Figure 4D, Video S4**). The activation of these integrins requires the same signaling molecules as did the activation of integrins in response to bead-based mimics (**Figure 4E)**. While these data show that MERTK signalling activates integrins and coordinates synapse formation, it is unclear how the spatial separation of MERTK and integrins occurs.

**Figure 4:**
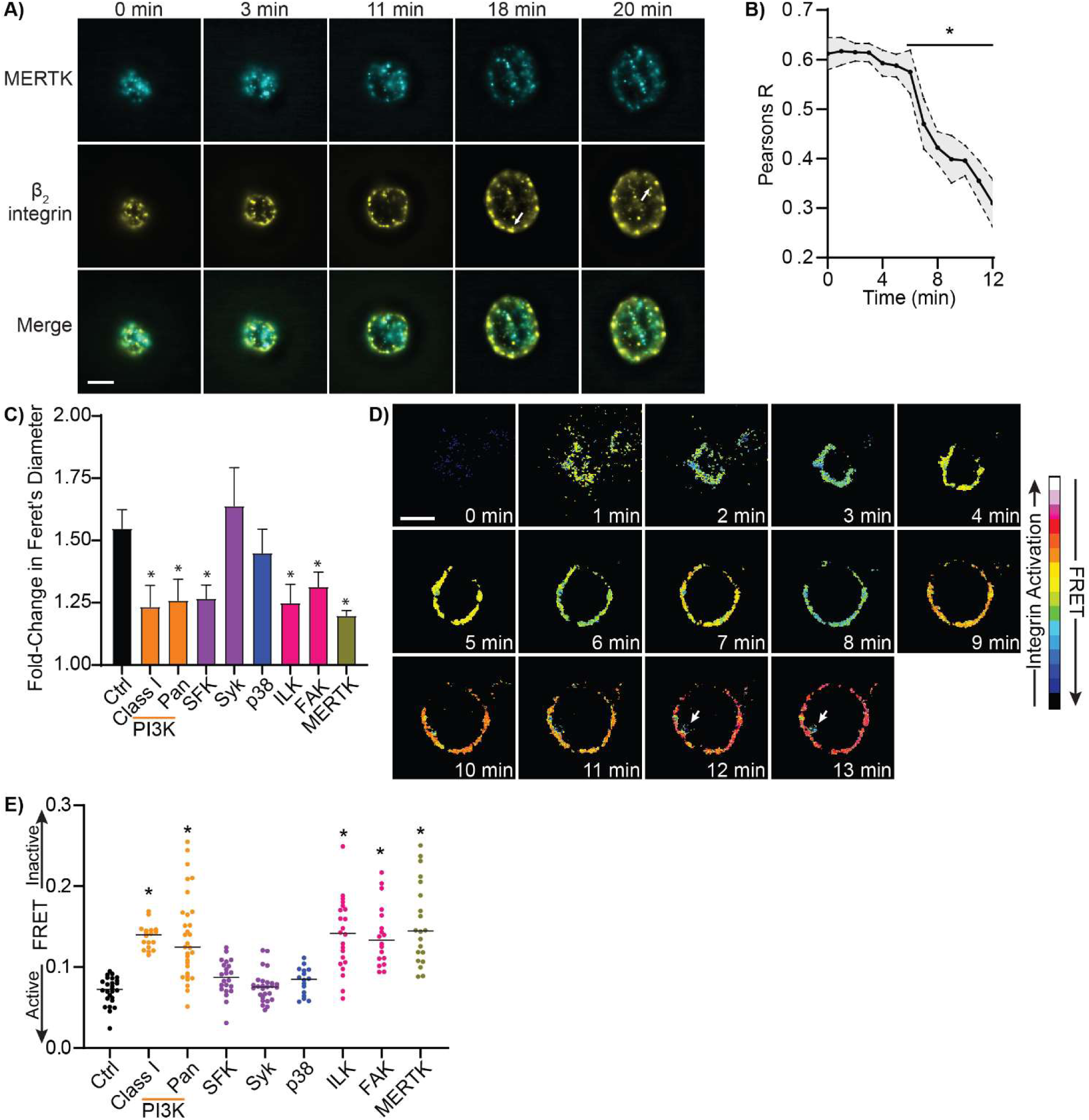
MERTK Signaling is Required for Efferocytic Synapse Formation. **A)** Timelapse micrograph showing the localization of MERTK (cyan) and β_2_ integrin (yellow) during the formation of an efferocytic synapse on a planar apoptotic cell mimic. Arrows indicate sites where β_2_ was exocytosed within the formed synapse. **B)** Quantification of MERTK co-localization with β_2_ integrin during efferocytic synapse formation on a planar apoptotic cell mimic. **C)** Quantification of the effect of PI3K, SFKs, Syk, p38 MAPK, FAK, ILK, MERTK kinase domain inhibition on the formation of an efferocytic synapse. Data is quantified in the fold-change of the maximum Feret’s diameter of the synapse compared to t = 0. **D)** Activation of β_2_ integrin in a developing efferocytic synapse, as quantified by FRET microscopy. Arrows indicate sites where β_2_ was exocytosed within the formed synapse. **E)** Effect of inhibitors of PI3K (LY & Wort), SFK (PP1), Syk (SykI), p38 MAPK (SB), ILK (ILK-I), FAK (PF), MERTK (UNC), and vehicle control on β_2_ integrin activation on opsonized planar apoptotic mimics containing 20:80 PtdSer:PtdChol. Data is plotted as mean ± SEM (B,C), as individual synapses (E), or as representative micrographs of individual cells (A,D), 0 min is the time when the cell fist enters the same focal plane as the planar apoptotic cell mimic. * = p < 0.05 compared to 0 min (B) or UT (C,E), Kruskal-Wallis test with Dunn’s multiple comparisons test.

### Diffusion Shapes the Efferocytic Synapse

There must be a difference in the movement of MERTK and β_2_ integrin within the synapse to account for their separation into separate spatial domains. Proteins in the plasma membrane diffuse via two distinct diffusive patterns: freely diffusing proteins which diffuse in a pattern that approximates Brownian diffusion, and proteins whose diffusion is confined to small sub-regions of the plasma membrane [46]. These confinement zones are created by “picket-fences” comprised of membrane-contacting actin fibril “fences” and actin-anchored transmembrane protein “pickets”. Combined, these create structures termed “corrals” which can confine protein diffusion to sub-region of the membrane over moderate (seconds-to-minutes) periods of time [47]. Given that active integrins bind to actin via talin [48], and our immunoprecipitation of MERTK did not identify any actin-binding proteins, changes in the actin cytoskeleton may serve to both form the synapse and to separate MERTK from integrins. The distribution of actin was imaged at 100 nm resolution using SRRF microscopy in macrophages forming synapses on PtdSer-containing planar bilayers. Actin was found in a relatively even meshwork beneath the plasma membrane before synapse formation, but following contact with the bilayer actin was readily redistributed to a ring at the edge of the synapse (**Figure 5A-C, Videos S5**). This change in actin distribution was dependent on the presence of PtdSer and was not observed on PtdChol bilayers (**Figure S5D, Video S6**). As expected, MERTK remained distributed throughout the synapse during synapse formation, while β_2_ integrin was moved to the outer edge (**Figure 5B-C**).

**Figure 5:**
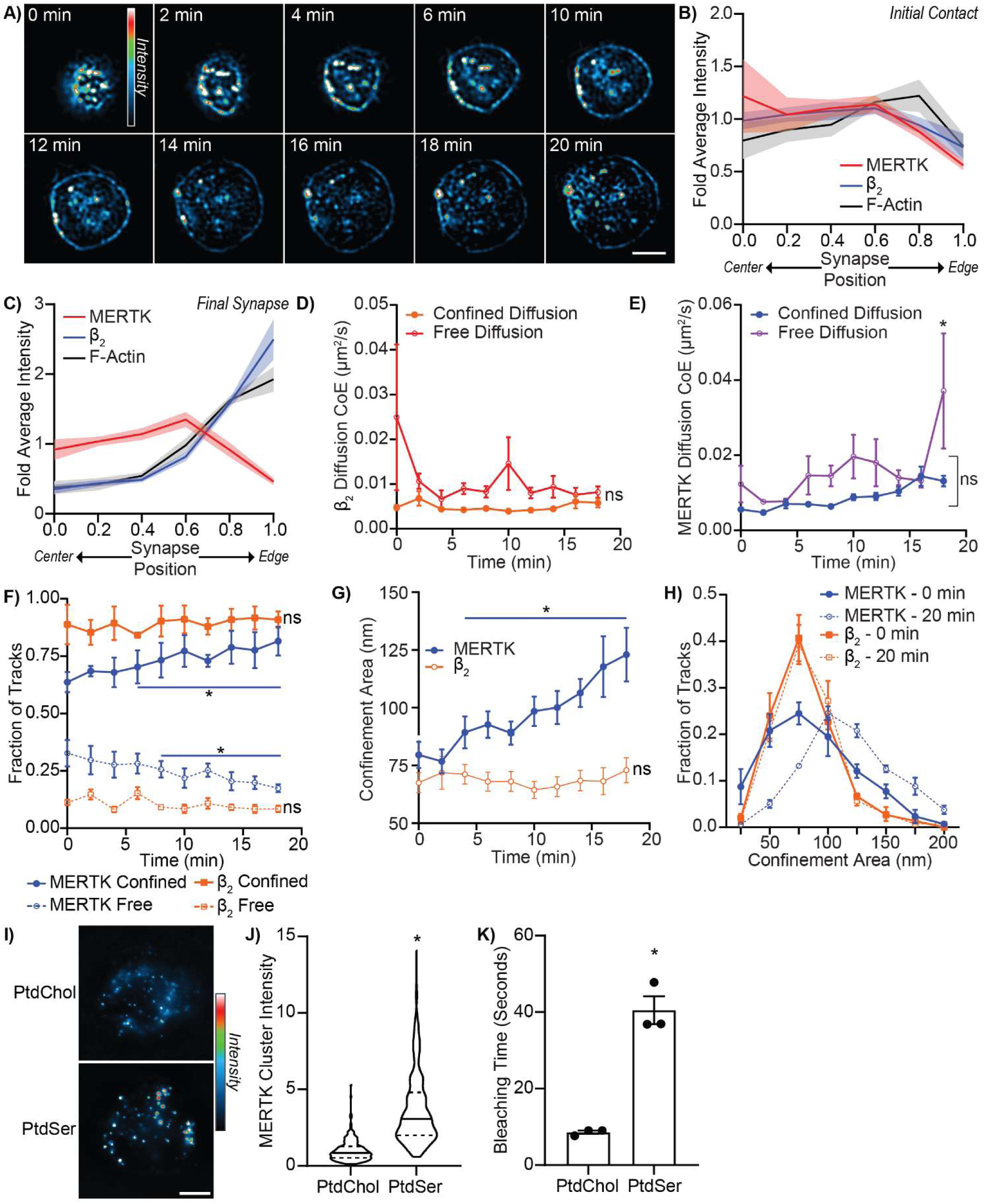
Changes in Diffusion Structures the Efferocytic Synapse. Efferocytic synapse formation was quantified on apoptotic cell-mimicking planar lipid bilayers comprised of 20% PtdSer and 80% PtdChol (PtdSer) or on bilayers mimicking a healthy cell (100% PtdChol). **A)** SRRF image of actin distribution during efferocytic synapse formation on a PtdSer-containing bilayer. **B-C)** Radial distribution of MERTK, β_2_ integrin, and F-actin in macrophages immediately after contacting a PtdSer-containing bilayer (B) and once efferocytic synapse formation was complete (C, 12 minutes post-contact). D-E) Changes in the diffusion rate of β_2_ integrins (D) and MERTK (E) during efferocytic synapse formation on a PtdSer-containing bilayer. Diffusion has been classified as either free diffusion or confined diffusion using moment scaling spectrum analysis. F) Quantification of the portion of MERTK and β_2_ integrin undergoing free diffusion (Free) versus confined diffusion during efferocytic synapse formation on a PtdSer-containing bilayer. G) Changes in the size of the confinement areas restricting the diffusion of MERTK and β_2_ integrin during efferocytic synapse formation on a PtdSer-containing bilayer. H) Size distribution of MERTK and β_2_ integrin confinement zones before (0 min) and after (20 min) efferocytic synapse formation on a PtdSer-containing bilayer. I-K) Images (I), integrated fluorescence intensity of individual clusters (J), and time to photobleach (K) of MERTK clusters on macrophages 20 minutes after contacting PtdChol versus PtdSer-containing bilayers. Data is presented as mean ± SEM, n = minimum of 5. 0 min is when the cell fist enters the focal plane. * p < 0.05 compared to t = 0, ANOVA with Tukey correction (E-G) or Mann Whitney U test (J-K). Scale bars are 10 μm.

Given the distribution of actin in the synapse, and actin’s role in confining the diffusion of transmembrane proteins, we expected to observe confined diffusion of β_2_ integrin at the synapse edge and highly diffusive MERTK in the synapse center [47,49]. Consistent with this model, diffusion analysis by single particle tracking microscopy revealed that β_2_ integrins had low diffusivity at all time points, while MERTK’s diffusivity increased as synapse formation progressed (**Figure 5D-E**). As expected, the large majority of β_2_ integrins underwent confined diffusion during synapse formation, with both the proportion of β_2_ integrins undergoing confined diffusion, and the size of the confinement areas, remaining consistent throughout synapse formation (**Figure 5F-G**). This pattern of high β_2_ integrin confinement and small confinement zone size is consistent with the integrins remaining bound to actin throughout synapse formation. Paradoxically, despite being localized to an area of the synapse which undergoes actin depletion during synapse formation, and exhibiting the increase in diffusivity expected to occur in areas of low actin density, the portion of MERTK undergoing confined diffusion increased as synapse formation progressed (**Figure 5F**). This increase in confined diffusion was accompanied by an increase in the size of the MERTK confinement zones (**Figure 5G)**. This increase in confinement region size occurred monotonically across all MERTK molecules in the synapse, indicating a change in the mechanism that confines MERTK diffusion rather than the appearance of a second MERTK population confined by a different mechanism (**Figure 5H**). On substrates lacking PtdSer, β_2_ integrin and MERTK exhibited no change in their diffusivity, fraction of free versus confined molecules, or confinement zone size (**Supplemental Figure 5E-G**), confirming that the changes in diffusion we observed on PtdSer-containing substrates was due to efferocytic signaling and not due to contact-dependent cellular responses or spatial crowding effects following cell-substrate contact.

Given the increase in MERTK confinement zone size, and the increase in MERTK diffusivity within these confinement zones, it seemed likely that the increase in MERTK diffusional confinement was due either to ligand-mediated immobilization of MERTK or due to clustering-induced immobilization. The former is unlikely given that apical-face lipids in supported lipid bilayers are highly diffusive and experience little diffusional confinement [50], while the latter is frequently observed [51,52]. To test whether MERTK was forming larger clusters within the synapse, we used stepwise photobleaching to measure the relative number of MERTK present in MERTK clusters on cells contacting PtdSer-containing versus PtdSer-free planar bilayers (**Figure 5I**). Consistent with ligand-induced receptor clustering driving the immobilization of MERTK in the synapse, MERTK clusters increased over four times in size on PtdSer-containing planar bilayers, as measured by both the integrated fluorescent intensity of individual clusters prior to photobleaching (**Figure 5J**) and by the time it took to completely photobleach these clusters at full excitation intensity (**Figure 5K**). Combined, these data support a model wherein β_2_ integrins are moved to the leading edge of the forming efferocytic synapse via their physical linkage to the expanding ring of actin that forms the synapse edge, while MERTK is retained in the centre of the synapse by ligand-induced clustering and the resulting immobilization of the receptor-ligand clusters.

## Discussion

While nearly a dozen efferocytic receptors are expressed by macrophages, MERTK is the predominant efferocytic receptor used by these cells, and is the primary or sole efferocytic receptor in multiple tissues including the heart and eyes [8,53–55]. SNPs in MERTK and its opsonins predispose individuals to a range of inflammatory and autoimmune disorders, highlighting the importance of this receptor to human health [30,31,56]. In this study we have demonstrated that MERTK activates a PI3K/SFK/ILK/FAK-dependent signaling pathway via its intrinsic kinase domain. The resulting PI3K activity induces the inside-out activation of β_2_ integrins, which then bind to apoptotic cells and mediate the expansion of an efferocytic synapse via an SFK-, ILK-, and FAK-dependent pathway. The differential diffusion of MERTK and β_2_ integrins is required for proper structuring of this synapse, with the diffusional confinement of β_2_ integrins retaining them at the actin-rich edge of the synapse, and ligand-induced clustering retaining MERTK in the actin-poor synapse centre.

The efferocytic synapse formed by MERTK is reminiscent of the phagocytic synapse formed during antibody-mediated phagocytosis by FcγRs. This phagocytic synapse forms a similar actin ring, which in early phagocytosis serves to expand the synapse, and later in phagocytosis serves as a “jaw” that exerts constrictive forces on the phagocytic target that drive phagosome closure [22,57–59]. The efferocytic and phagocytic synapses also contain a similar bounding ring of integrins, which during IgG mediated phagocytosis serves to exclude proteins with a large glycocalyx from the phagocytic synapse [23]. This process excludes inhibitory phosphatases such as CD45 and CD148 in a manner similar to the exclusion of phosphatases from the immunological (T cell) synapse [60]. But while the efferocytic synapse shares a similar bounding ring of integrins, it is unlikely that these integrins are playing the same role. Indeed, at the time of this writing there are no published reports showing a role for CD45 or CD148 in the inhibition of efferocytosis. Moreover, phagocytosis needs to be sensitive to the ligation of a small number of receptors in order to ensure the removal of poorly opsonized or small pathogens, with the exclusion of phosphatases from small receptor clusters allowing for the unopposed activation of the phagocytic signalling pathway [23]. In contrast, efferocytosis needs to be tightly regulated, as the “eat me” signal PtdSer is frequently exposed on non-apoptotic cells, including on platelets during coagulation, on degranulating mast cells, on migratory CD45RB^lo^ T cells, on CD8^+^ T cells following antigen recognition, in skeletal muscle, on activated and IL-10 producing B cells, and following galectin binding to neutrophils [61–69]. Thus, formation of the efferocytic synapse must be highly regulated to prevent engulfment of PtdSer-positive non-apoptotic cells. This regulation is mediated by the “don’t eat me” receptor CD47 on non-apoptotic cells and its cognate receptor SIRPα on the efferocyte [70]. Unlike CD45 and CD148, neither CD47 or SIRPα are excluded from the efferocytic synapse, thus ensuring that their inhibitory functions are retained even when efferocytic ligands are present [71]. Instead of relying on exclusion, the inhibitory signaling of SIRPα is regulated via CD47 clustering, with apoptosis-induced dispersion of CD47 clusters eliminating this inhibitory signal and thus allowing efferocytosis to occur [72].

Although integrins are not acting as an exclusionary barrier for CD47/SIRPα in the efferocytic synapse, they do serve a critical role in efferocytosis as our data indicate that MERTK is unable to mediate the engulfment of apoptotic cells in the absence of integrin ligands, or when MERTK-mediated integrin activation is inhibited [71]. While conjectural, we believe that MERTK is utilizing integrins to mediate high-affinity binding and actin reorganization during efferocytosis. Indeed, the equilibrium dissociation constant of integrin-ligand pairs is typically in the range of 10^-9^ to 10^-10^ M, which is one-to two-orders of magnitude higher than the affinity of MERTK and other TAMs for Gas6 and Protein S, with integrin clustering further increasing the avidity of this interaction [73–76]. Moreover, integrins can independently mediate the actin reorganization processes required for engulfment of large particulate targets, most notably in the form of complement-mediated phagocytosis where the entirety of the phagocytic process is mediated by β_2_ integrin-iC3b interactions and integrin signaling (reviewed in [42]). Further evidence that MERTK is utilizing the integrin-mediated phagocytosis pathway can be found in the signaling molecules required for MERTK-mediated efferocytosis. Herein, we determined that the kinase Syk was not required for MERTK-mediated efferocytosis, while ILK and FAK were essential. Complement-mediated phagocytosis displays the same independence from Syk signaling, whereas Syk is required for antibody-mediated phagocytosis [77,78]. Likewise, FAK signaling is inhibitory for antibody mediated phagocytosis, whereas both FAK and ILK are required for integrin-mediated phagocytosis [79–81]. Critically, we were able to induce the engulfment of sCD93-opsonized (e.g. α_x_β_2_ integrin-specific) efferocytic targets simply by forcing integrin activation by the addition of manganese, and we were able to abrogate integrin activation in the presence of opsonized efferocytic targets through inhibiting MERTK kinase activity or by knocking-down MERTK. Consistent with efferocytic receptors activating integrins in order to induce engulfment, ligated SIRPα inhibits efferocytosis by preventing integrin activation [71]. Combined, these data indicate that MERTK acts as a sentinel receptor which identifies apoptotic cells and then activates integrins which then engage the canonical integrin-mediated phagocytosis pathway to drive the engulfment of the apoptotic cell.

Previous studies analyzing other efferocytic receptors have found a similar dependence on integrins, leading to the tether model which proposed that efferocytic receptors such as MERTK act as passive “tethers” that stabilize the association of apoptotic cells with efferocytes, thereby providing integrins the time necessary to recognize and bind to the apoptotic cell [7,82]. Our data challenges this model, and indeed, demonstrates that MERTK generates a signal via its kinase domain which activates PI3K, which in-turn activates integrins and allows efferocytosis to proceed. Without the activating signals from the MERTK kinase domain and PI3K, neither integrin activation nor engulfment of apoptotic targets occurs. While this is not the first indication that efferocytic receptors are more than passive receptors [14], to our knowledge our data is the first identifying a specific signaling pathway induced by an efferocytic receptor that then regulates integrin affinity and activation. Interestingly, multiple other efferocytic receptors are known or suspected to associate with MERTK or another member of the TAM family – e.g. TIM4 and BAI1 with MERTK, and TIM1 with Tyro3 [83–85], with these interactions often having a synergistic effect on efferocytosis. Neither the TIM family of receptors nor BAI1 have intrinsic kinase domains, suggesting that they may act as classical tethers and/or require the kinase activity of TAM family kinases to initiate signaling. Indeed, while BAI1 was found to interact with the ELMO/DOCK180 complex that initiates the actin reorganization required for phagocytosis/efferocytosis, this complex’s activity is initiated by a tyrosine-kinase mediated phosphorylation of CrkII, with TAMs known to be capable of providing this activating phosphorylation [35,86,87]. While our mass spectrometry analysis did not identify any proteins known to act as intermediaries between MERTK and PI3K, MERTK is known to directly phosphorylate and activate the PI3K effector AKT, and immunoprecipitations using a soluble MERTK kinase domain as bait identified the scaffolding protein Grb2 as a major interactor [33,88]. Grb2 is known to act as a bridge between phosphorylated receptor tyrosine kinases and PI3K, and indeed, Grb2 has been shown to bind to the phosphorylated (active) MERTK kinase domain where it mediates recruitment and activation of type 1A PI3-kinases through interactions with the SH2 domain in the p85 subunit of PI3K [89]. While most studies identify PI3K signaling as a product of outside-in signaling through integrins, our observation of PI3K-mediated inside-out activation is not unprecedented. Thamilselvan *et al.* identified a pressure-induced pathway resulting in the PI3K-dependent activation of β_1_ integrins, while both CD44 mediated phagocytosis and LFA-1 mediated T cell adhesion occur via the PI3K-dependent activation of the GTPase Rap-1, which then activates β_2_ integrins [90,91].

In summary, we have identified the sequential series of events which allows MERTK to activate integrins and mediate the formation of a functional efferocytic synapse. MERTK ligation induces activation of its kinase domain which, via Grb2, activates PI3K [33]. PI3K activity then induces the inside-out activation of integrins that then mediate the actin reorganization needed for expansion of the synapse around the apoptotic cell in a SFK, ILK, and FAK-dependent manner. Given the structural similarities of MERTK with the other TAM family members Axl and Tyro3, this mechanism is likely universal across this family of efferocytic receptors.

## Materials & Methods

### Materials

THP-1 cells were obtained from Cedar Lane Labs (Mississauga, Canada). Roswell Park Memorial Institute (RPMI), Dulbecco’s Modified Eagle Medium (DMEM), and Fetal Bovine Serum (FBS) were purchased from Wisent (Saint-Jean-Baptiste, Canada), Trypsin-EDTA and antibiotic/antimycotic were purchased from Corning (Manassas, Virginia). #1.5 thickness 18 mm round cover slips and 16% paraformaldehyde (PFA) were purchased from Electron Microscopy Supplies (Hatfield, Pennsylvania). Anti-Syk clone EP573Y and anti-FAK clone EP695Y were from Abcam (Cambridge, United Kingdom). Anti-MERTK clone LS-C199253 was from LSBio (Seattle WA). CD18 clone TS1/18, Octadecyl rhodamine B, Hoechst, Permafluor mounting medium, N-hydroxysuccinimidobiotin and cell proliferation dye 670 were purchased from Thermo Fisher Scientific. Silica beads were purchased from Bangs Laboratories, Inc. (Fishers, Indiana). Lipids, biotin PE, and the liposome extruder were from Avanti Polar Lipids (Alabaster, Alabama). Gas6 was purchased from R&D (Minneapolis, MN). FITC anti-human CD18 antibody and FITC anti-human CD11c antibody (clone Bul15) were purchased from BioLegend (San Diego, USA). Rabbit anti-human MERTK antibody (D21F11) was purchased from Cell Signaling Technology (Danvers, Massachusetts). Fluorescent secondary antibodies were purchased from Developmental Studies Hybridoma Bank (Iowa City, Iowa), and Jackson Immuno Research Laboratories, Inc. (West Grove, Pennsylvania). ILK inhibitor Cpd-22 was purchased from EMD Millipore. The FAK inhibitor PF-00562271 was purchased from Sigma-Aldrich, and the other pharmacological inhibitors for Src-family kinases (PP1), Syk (Syk Inhibitor I), class-I PI3K (Ly294002), pan-PI3K (Wortmannin), and p38 MAPK (SB 203580) were purchased from Cayman Chemical. Phorbol 12-Myristate 13-Acetate (PMA) and rest of the chemicals were purchased from BioShop Canada (Mississauga, Canada). ON-TARGET siRNA’s against MERTK, Tyro3, Axl and αx integrins were purchased from Horizon Discovery (Cambridge, UK). Soluble CD93 was produced as described previously [16]. Partek Flow was from Parteck (St. Louis, MO) MATLAB and Prism software were purchased from MathWorks (Natick, Massachusetts) and GraphPad (La Jolla, California) respectively. FIJI was downloaded from https://fiji.sc/ [92].

### THP-1 Cell Culture and Differentiation

THP-1 cells were maintained in RPMI supplemented with 10% FBS at 37°C in 5% CO_2_. Cells were split upon reaching 1.5 ×10^6^ cells/mL. To differentiate THP-1 cells into macrophages, cells were collected by centrifugation and resuspended in fresh medium at 2.5 × 10^5^ cells/ml, and 100 ng/ml PMA added. These cells were then seeded into 12-well tissue culture plate with #1.5 thickness 18 mm diameter coverslips plated into each well at a density of 2.5 × 10^5^ cells/well and differentiated for 72 hours at 37°C in 5% CO_2_.

### Human Peripheral Blood Macrophage Differentiation

The collection of blood from healthy donors was approved by the Health Science Research Ethics Board of the University of Western Ontario and was performed in accordance with the guidelines of the Tri-Council policy statement on human research. Blood was drawn into heparinized vacuum collection tubes, layered on an equal volume of Lympholyte-poly and centrifuged at 300 × *g* for 35 min at 20 °C. The top band of peripheral blood mononuclear cells was collected, washed once (300 × *g*, 6 min, 20 °C) with PBS, resuspended in completed medium (RPMI-1640+10% FBS and 1% antibiotic–antimycotic solution), and ∼1 × 10^6^ cells placed into each well of a 12-well plate. The cells were allowed to adhere for 1 hr at 37°C/5% CO_2_, then the wells washed gently with PBS to remove non-adherent cells, and the cells cultured in complete medium with 10 ng/mL M-CSF (10 ng/ml) for 7 days.

### RNAseq

Expression of efferocytic receptors in THP-1 macrophages was determined by searching our previously published RNAseq dataset for known efferocytic receptors and opsonins [93]. Briefly, total RNA was isolated from PMA-differentiated THP-1 macrophages and processed with a Vazyme VAHTS Total RNA-seq Library Prep Kit for Illumina, single-end reads sequenced on an Illumina NextSeq 500, 1 x76 bp, using 75 cycles of a High Output v2 kit. Fastq data files were analyzed using Partek Flow (St. Louis, MO) and the sequences were aligned to the *Homo sapiens* genome using STAR 2.6.1d and annotated using Ensembl v 98. Gene expression levels were normalized Reads Per Kilobase of transcript, per Million mapped reads (RPKM), and genes with fewer than 10 reads removed from the dataset. This dataset is available at https://doi.org/10.5683/SP2/ISOA8W.

### Mass Spectrometry

HA-MERTK expressing cells were prepared previously [41] and cultured in DMEM + 10% FBS at 37°C/5% CO_2_. These cells were washed twice with PBS and then reversibly crosslinked using 0.5 mM Dithiobis[succinimidyl propionate] and 1.0 mM Dithio-bismaleimidoethane for 20 min at room temperature. Crosslinking was stopped by the addition of 1 ml of quenching buffer (5 mM l-cysteine+20 mM Tris-Cl, pH 7.4) for 10 min, then the cells lysed in 2 mL of RIPA buffer supplemented with 1 mM NaVO_4_, 0.25 mM PMSF, 200 nM okadaic acid, 10 mM NaF, and HALT protease inhibitor cocktail. 1 µg of anti-HA was added to half the cell lysate, 1 µg of an isotype control added to the other half, and both incubated for 2 hr at 4°C, at which point 20 µL of Protein A/G beads were added to each tube, and the lysates incubated an additional hour at 4°C. The beads were pelleted by a 1 min/16,000 × g centrifugation and washed three times with RIPA buffer using 1 min/16,000 × g centrifugation. After the final wash the beads were pelleted, resuspended in 50 µL of Laemmli buffer + 350 mM DTT, and heated at 98°C for 5 minutes. The beads were centrifuged at 21,000 × g for 5 mins and the supernatant was then loaded into a 4%-20% gradient SDS-PAGE gel, run at 100 V. The gel was fixed with 40% ethanol/10% acetic acid for 15 mins and then stained with colloidal Coomassie stain as per the manufacture’s instructions. Bands unique to the anti-HA lane were excised with an Ettan Robotic Spot- Picker, destained with 50% acetonitrile + 50 mM ammonium bicarbonate, treated with 10 mM DTT and 55 mM iodoacetamide, and digested with trypsin. Peptides were extracted with 1% formic acid + 2% acetonitrile, dried, and then suspended in water with 0.1% trifluoroacetic acid and 10% acetonitrile. Mass spectrometry was performed on an AB Sciex 5800 TOF/TOF System and analyzed using a MASCOT database search.

### Microscopy

Other than GSDM, all microscopy was performed on a Leica DMI6000B microscope equipped with a photometrics Evolve-512 delta EM-CCD camera, Chroma Sedat Quad filter set, heated/CO_2_ perfused live-cell stage, 100×/1.40 NA and 60×/1.40 NA objectives, adaptive focus control, and running Leica LAX image acquisition software. All live-cell imaging is performed in imaging buffer (150 mM NaCl, 5 mM KCl, 1 mM MgCl_2_, 100 µM EGTA, 2 mM CaCl_2_, 20 mM HEPES and 2 g/L sodium bicarbonate, pH 7.4).

### Ground State Depletion Microscopy

THP-1 macrophages were grown on coverslips as described above, and then fixed for 15 min at 37°C in PEM buffer (80 mM PIPES pH 6.8, 5 mM EGTA, 2 mM MgCl_2_) + 4% PFA to limit artificial receptor clustering [94]. For extracellular staining samples were then washed once with PBS and blocked for 1 hr at room temperature with PBS + 2.5% BSA, while for intracellular staining cells were washed once with PBS and blocked/permeabilized for 1 hour with PBS + 2.5% BSA + 0.1% Triton X-100. Next, samples were stained with anti-MERTK plus one or two of (depending on species and fluorophore compatibility): anti-β_2_ integrin, anti-α_x_ integrin, anti-Syk, or anti-FAK and incubated for 20 min (extracellular labeling) or 1 hr (intracellular labeling) followed by 3 × 15 min washes in PBS. Samples were then stained with fluorescently labeled Fab fragments, diluted 1:1,000 in PBS + 2.5% BSA, to avoid cross-linking [95,96], using the same incubation times and washes as used for primary antibodies. The cells were post-fixed with 2%PFA+PEM at room temperature for 20 minutes. Samples were immediately transferred to a chamber slide containing PBS + 100 mg/mL β-mercaptoethylamine and transferred to the stage of a Leica GSDM microscope. GSDM acquisitions of the basolateral cell surface were acquired in TIRF mode and the resulting molecular coordinate lists imported into our MIISR analysis software which was then used to perform SAA and Ripley’s clustering analysis [96].

### Preparation of Phagocytic and Efferocytic Targets

Mimics of antibody-opsonized pathogens were prepared by mixing 10 μL of 3 μm polystyrene-divinylbenzene beads with 100μL of PBS and 10μL human IgG, then rotating for 2 hours at room temperature. Beads were washed twice with 1 mL of PBS and a 4,500 × g/1 min centrifugation, then re-suspended with 100μL of PBS. Efferocytic targets were prepared by combining 10 μL of 3.17μm silica beads with 25 nmol of a 79.9:20:0.1 molar mixture of Phosphatidylcholine (PtdChol):PtdSer:biotin-Phosphatidylethanolamine (biotin-PE) in chloroform. Chloroform was then evaporated under nitrogen gas, the beads washed twice in 500 μL PBS by centrifuging at 4,500 x g for 1 min, and then resuspended in 100 μL PBS. Opsonized PtdSer/PtdChol beads were prepared by rotating 3 μL of the lipid-coated bead preparation with 3 μL of recombinant Gas6 and/or 5 μL of recombinant MFG-E8 in a total volume of 25µl PBS for 2 hours at room temperature and washed once with PBS. For sCD93-opsonized beads, 3 μm polystyrene/DVB beads were coated with 3 μL of recombinant Gas6 and/or 5 μL of recombinant sCD93 as above, and then any remaining protein binding sites blocked by addition of an equal volume of 1% BSA in PBS for 30 min.

### Apoptotic Cell Targets

Apoptotic cells were prepared as described previously [97]. Briefly, Jurkat T cells were grown in RPMI + 10% FBS, pelleted with a 500 × g/5 min centrifugation, suspended in serum-free medium containing 1 µM staurosporine at 2 × 10^6^ cells/mL, and cultured overnight (∼16 hr) at 37°C in 5% CO_2_. The cells were then pelleted, resuspended in 1 mL PBS + ∼0.005 mg N-hydroxysuccinimidobiotin and 5 µM cell proliferation dye 670, and incubated at room temperature for 20 min. 10% (v/v) human serum or an equal volume of 2.5% BSA in PBS was added to quench any unused dye. If BSA was used for quenching, the apoptotic cells were subsequently opsonized by addition of 3 μL of recombinant Gas6 and/or 5 μL of recombinant sCD93 for 2 hours at room temperature, and washed once with PBS. Apoptotic cells were used immediately after preparation.

### Inhibitor and siRNA Treatment

For assays using pharmacological inhibitors, cells were pre-treated with 5 µM of the MERTk kinase domain inhibitor UNC2250 [40], 2.5 μM of the class I PI3K inhibitor LY294002, 100 nM of the pan-PI3IK inhibitor Wortmannin, 30 μM of the SFK inhibitor PP1, 100 nM Syk inhibitor I, 10 µM of the p38 MAPK inhibitor SB 203580, 2 µM of the ILK inhibitor Cpd-22 inhibitor, or 25 µM of the FAK inhibitor PF-00562271, for 15 minutes at 37 °C prior to starting the experiment, and the inhibitors maintained at these concentrations throughout the experiment [98,99]. For assays using siRNA, THP-1 macrophages were suspended by scraping, collected by a 600 × g, 5 min centrifugation, and resuspended at 2 × 10^7^/ml in buffer R, to which 150 μM of siRNA was added and the sample electroporated using a Neon electroporation system with 2 × 20 ms pulses at 1400 V. The cells were then placed onto glass coverslips placed in a 12-well plate, along with 1 ml of RPMI + 10% FBS, and allowed to recover for a minimum of 24 hr in a 37°C/5% CO_2_ incubator. Knockdown was confirmed by semi-quantitative RT-PCR. Briefly, total RNA was purified and reverse transcribed, amplified using multiplex PCR primers for MERTK (5’-CCA GGT GAC CTC TGT CGA ATC AAA-3’, 5’-GGG ATT TTT CAG GCT GTT CGT TAA CA-3’), Axl (5’-CTC CAC CTG GTC TCC CG-3’, 5’-CCA GGG GCA CTC CCT-3’), Tyro3 (5’-CTC TTC TCA TGA CCG TGC AGG-3’, 5’-CCG GGC CAT GAC ACT GT-3’), α_x_ integrin (5’-GGT TTC AAC TTG GAC ACA GAG GAG C-3’, 5’-GAAGGG CTG GTG GTA GAC GC-3’), and resolved using a 1.5% agarose gel.

### Phagocytosis and Efferocytosis Assays

Phagocytosis and efferocytosis assays were performed as described previously [97,100]. Phagocytosis/efferocytosis was quantified by adding 2 µL of IgG or lipid coated beads per well of macrophages (∼200,000 beads/well, or 800 beads/mm^2^), or adding 5 × 10^5^ apoptotic cells/well. Cells were centrifuged at 400 × g for 1 minute to force beads into contact with macrophages, then incubated at 37°C with 5% CO_2_ for 60 minutes to allow for phagocytosis/efferocytosis to occur. The cells were then washed three times with cold PBS. Non-internalized IgG beads were labeled using 1:500 dilution of goat anti-human Alexa488 antibody, non-internalized lipid coated beads and apoptotic cells were labeled with 1:500 fluorescent streptavidin, for 20 minutes. Cells were then washed three times with PBS, fixed for 15 min with 4% PFA in PBS, and mounted in a Leiden chamber for imaging. For quantification, phagocytic and efferocytic index were calculated using beads as the average number of beads internalized per macrophage, while the Alexa488/Alexa647 was used to identify the number of bound beads per macrophage. For apoptotic cell uptake, an Otsu threshold was applied to the streptavidin image, the resulting mask inverted, and the inverted mask applied to the cell proliferation dye 670 image to eliminate signal from non-internalized apoptotic cells. The DIC image was then used to manually create polygonal ROIs around each macrophage, and the integrated cell proliferation dye 670 signal measured for each ROI. The integrated intensity of each cell was then normalized to the average integrated intensity of the control cells, with these averaged values used for comparisons between experimental repeats.

### Immunostaining

THP-1 cells were washed with PBS and fixed with 4% PFA in PBS at 10°C for 20 minutes. Cells were washed three times with PBS, followed by 1 hr incubation with primary antibody (1:200, anti-human MERTK, 1:40, FITC anti-human CD18, anti-human CD11c and anti-human CD11b monoclonal antibodies). Cells were then washed with 3 × 15 min with PBS and stained using 1:200 fluorescently tagged secondary antibodies. Cells were washed 3 × 15 min with PBS, incubated for 5 min with 1:10,000 Hoechst in PBS for 5 mins, and washed a final time in PBS before being mounted on a glass slides with PermaFluor.

### Integrin Activation FRET Assay

THP-1 cells were plated and differentiated as described above, then cooled to 10°C for 10 min. Cells were washed 3 times with PBS and incubated with a FITC anti-human CD18 antibody (0.25 µg/ml) for 20 minutes at 10°C. After 20 minutes, cells were washed with PBS and incubated with 100 nM octadecyl-rhodamine B (ORB). Unstained, donor-only (FITC-stained) and acceptor-only (ORB-stained) samples were also prepared. Cells were then washed with PBS and pharmacological inhibitors added at the concentrations indicated previously. The coverslip was then transferred to a Leiden chamber, and filled with imaging buffer containing Gas6-opsonized apoptotic cell mimics (800 beads/mm^2^), and the chamber transferred to the pre-heated stage of the Leica DMI6000B microscope and imaged for donor fluorescence (Idd—λEx/λEm: 488 nm/525 nm), FRET (Ida—λEx/λEm: 488 nm/578 nm), acceptor fluorescence (Iaa—λEx/λEm: 555 nm/575 nm) and in DIC. As a positive control, 1 mM of Mn^2+^ was added to some samples to force integrins into their active conformation. Images were captured every 2 min for up to an hour. The resulting images were imported into FIJI and corrected sensitized emission FRET calculated using the method of van Rheenen *et al* [101]. Briefly, unstained cells were used to quantify the cellular background in the Idd, Ida and Iaa channels, and this value subtracted from the corresponding channel in all images. Then, donor cross-talk (β, donor-only Ida/Idd), donor cross-excitation (α, acceptor-only Idd/Iaa), acceptor cross-excitation (γ, acceptor-only Ida/Iaa), and FRET cross-talk (δ, acceptor-only Idd/Ida), were calculated. FRET efficiency (E_A_) was then calculated in background subtracted images using the formula:

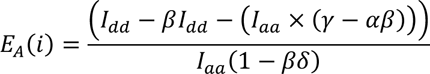

Lastly, a mask was generated by applying an Otsu threshold to the Idd channel, which was then dilated once, and the resulting mask applied to the FRET image to eliminate spurious signals from regions lacking β_2_ integrins. This 32-bit image was used to quantify integrin activation, but for images displaying the localization of integrin activation, this image was converted to a 16-bit image, inverted to display integrin activation rather than raw FRET signal, and a 16-color ramp LUT applied.

### Supported Lipid Bilayer Preparation

#1.5 thickness, 18 mm diameter glass coverslips were cleaned in 70% ethanol, distilled water, and then heated to 55 °C in 2 M HCl to remove all organic material. These coverslips were then plasma cleaned by a 10 second exposure to air-plasma (<1,000 mTorr) in a homemade 1500 W microwave plasma chamber [102]. 0.02 µmol of 100 μm diameter liposomes mimicking either a healthy cell (99.9:0.1 molar ratio of PtdChol:Biotin-PtdEth) or apoptotic cell (79.9:20:0.1 molar ratio of PtdChol:PtdSer:biotin-PtdEth) were prepared as per the manufacture’s instructions and placed on a plasma cleaned coverslip to self-deposit into a supported lipid bilayer (SLB). For some experiments the biotin-PtdEth was excluded from the lipid mixture, and instead, a small pen mark was placed on the top surface of the coverslip prior to addition of the liposomes. Coverslips were incubated for 25 min, then washed 3 × with PBS, then incubated for 25 min with 2.5% BSA in PBS ± 1:1000 strepatavadin-alexa-647 to block any non-lipidated surfaces and to label the monolayer. The SLB’s were then washed 3× with PBS and placed in a Leiden chamber filled with imaging buffer.

### Efferocytic Synapse Imaging

∼2.5 × 10^5^ PMA-differentiated THP-1 macrophages were cooled to 10 °C, and washed 3× with PBS. For FRET imaging, cells were labeled as described above. For synapse and focal exocytosis imaging the cells were labeled with 0.25 μg/mL FITC-conjugated mouse-anti-human CD18 antibody plus 0.25 μg/mL rabbit-anti-human-MERTK for 20 min at 10°C, washed 3 × with PBS and then labeled with a Cy3-labeled anti-rabbit secondary Fab. For single-particle tracking (SPT), cells were labeled as for synapse imaging, but using a goat-anti-rabbit Alexa-647 secondary Fab in lieu of the Cy3-labeled Fab, producing a label density of ∼1 Cy3/Alexa647 per 2.5 μm^2^. After labeling the cells were washed 3 × with PBS and suspended by removing all medium and adding 300 µL of accutase for 10 min at room temperature. 700 μL of FBS-supplemented RPMI were added to each well to suspend the cells, the cell suspension pelleted with a 400 × g/4 min centrifugation, the supernatant was removed, the pellet was resuspended in 800 μL of imaging buffer + 10% human serum, and the cells added onto Leiden chamber containing an SLB-coated coverslip. For FRET and synapse/exocytosis imaging, 1.6 μm thick z-stacks (0.4 μm spacing between slices) were captured of the cells every minute, using the strepatavadin-Alexa647 labeled SLB to demark the lower bound of the z-stack, and capturing either Idd, Ida, Iaa and DIC channels (FRET imaging) or FITC, Cy3, Alexa647 and DIC imaging (synapse/exocytosis imaging). For SPT imaging, the position of the SLB was identified using the pen mark on the coverslip prior to addition of the cell. Every 2 minutes a single DIC, Alexa405 and FITC image was collected, as well as 50 frames at 100 ms/frame of the Cy3 and Alexa647 channels for SPT analysis. Integrin activation was quantified using FRET as described above. Colocalization between synapse components was quantified using Pearson correlation as implemented in the Just Another Colocalization Plugin in ImageJ/FIJI [92,103]. Synapse expansion was quantified by manually tracing the outer edge of the β_2_ ring using the polygon selection tool in ImageJ/FIJI and measuring the Ferrets diameter. This was repeated for cell at each time point, normalizing all diameters to the diameter of the synapse at the first time point a synapse was visible in the time-series. Focal exocytosis was identified using our previously published exocytosis algorithm [104].

### Single-Particle Tracking

Single particle tracking data was analyzed by exporting as TIFF-stacks the time-series of the Cy3 and Alexa647 acquisitions. Diffusion of MERTK and β_2_ integrin were quantified using the SPT-acquisitions as describe previously [105,106]. Briefly, SPT was performed using the Matlab scripts of Jaqaman *et al*., and custom-written Matlab scripts used to remove any molecular trajectories with a positional accuracy worse than 25 nm [107]. Moment scaling spectrum (MSS) analysis was used to classify the diffusion of MERTK and β_2_ [108], where MSS classifies molecular movement based on the power-law indices, γ_ρ_, of the first through fourth moments, ζ_K_^pr^〉, of molecular movement, assuming each moment has a power-law dependency on the lag-time (τ) of ζ_K_^pr^〉^yp^, where Brownian motion is defined as ρ ≈ y_ρ_, subdiffusive (confined) movement as ρ < y_ρ_, and superdiffusive movement as ρ > y_ρ_. The diffusion rate is estimated from the initial slope of mean-squared displacement (MSD) plot of each particle, and for confined particles, the confinement zone diameter is calculated as the 90^th^ percentile of the maximum possible extent of the particles variance-covariance matrix.

### MERTK Cluster Size Analysis

MERTK on THP-1-derived macrophages was labeled as described above, and the macrophages placed onto supported lipid bilayers containing either 100% PtdChol or 80% PtdChol + 20% PtdSer. 30 minutes later, the cells were imaged at 100 ms and maximum excitation energy (∼2 W/m^2^). MERTK cluster size was then quantified from this time series using two analyses. First, integrated intensity of MERTK clusters was quantified in the first image of these time-series using FIJI, by drawing an ROI around each stationary MERTK cluster, and the integrated intensity of each cluster normalized to the mean intensity of MERTK clusters on PtdChol-only substrates. As a second measure of cluster size, the time to photobleach the clusters was quantified from the time series, with complete photobleaching defined as a fluorescent intensity within 1 SD of the background fluorescence.

### Statistical Analysis

Unless otherwise indicated data are presented as the mean ± SEM and analyzed using a two-tailed Student’s T-test or ANOVA with Tukey correction. All statistical analyses were performed in GraphPad Prism (GraphPad, San Diego, CA).

## Acknowledgments & Funding

This work was funded by a Canadian Institutes of Health Research Project Grant (PJT-162203) to BH. BHD and ENB were funded by Canada Graduate Scholarships – Master’s (CGSM) from the CIHR. The funding agencies had no role in study design, data collection and analysis, decision to publish, or preparation of the manuscript.

## Supplemental Video Captions

**Supplemental Video 1.** Integrin Activation During Efferocytosis of Gas6-opsonized apoptotic cell mimics. DIC (top-left), TS1/18-FITC labeled of β_2_ integrins (top-right/yellow), ORB-labeled plasma membrane (lower-left/magenta), and integrin activation as measured by the inverse acceptor-sensitized FRET emission (lower-right, colour-coded) images are shown. Mimics engaged by the macrophage are indicated by white arrows. Scale bar is 10 μm. File available as Supplemental Video 1.mp4.

**Supplemental Video 2.** Localization of MERTK and of β_2_ integrin on a THP-1 macrophage forming a frustrated efferocytic synapse on a 20% PtdSer/80% PtdChol planar lipid bilayer. The cell is labeled with antibodies against MERTK and β_2_ integrin. Scale bar is 10 μm. File available as Supplemental Video 2.mp4.

**Supplemental Video 3.** β_2_ integrin localization, relative to the macrophage cell body. Scale bar is 10 μm. File available as Supplemental Video 3.mp4.

**Supplemental Video 4.** β_2_ integrin activation, as measured by FRET, during frustrated efferocytosis. Scale bar is 10 μm. File available as Supplemental Video 4.mp4.

**Supplemental Video 5.** Actin redistribution during efferocytic synapse formation within a frustrated efferocytosis assay. Scale bar is 10 μm File available as Supplemental Video 5.mp4.

**Supplemental Video 6.** Actin redistribution during efferocytic synapse formation on a planar bilayer comprised of PhtChol. Scale bar is 10 μm File available as Supplemental Video 6.mp4.

## Preprint Notice

This research article is a preprint and has not been peer reviewed.

## References

1. Brown GD, Gordon S. Immune recognition. A new receptor for beta-glucans. Nature. 2001;413: 36–37. doi:10.1038/35092620

2. Xu S, Wang J, Wang J-H, Springer TA. Distinct recognition of complement iC3b by integrins αXβ2 and αMβ2. Proc Natl Acad Sci U S A. 2017;114: 3403– 3408. doi:10.1073/pnas.1620881114

3. Bruhns P, Iannascoli B, England P, Mancardi D a, Fernandez N, Jorieux S, et al. Specificity and affinity of human Fcgamma receptors and their polymorphic variants for human IgG subclasses. Blood. 2009;113: 3716–3725. doi:10.1182/blood-2008-09-179754

4. Lam AL, Heit B. Having an Old Friend for Dinner: The Interplay between Apoptotic Cells and Efferocytes. Cells. 2021;10: 1265. doi:10.3390/cells10051265

5. Suzuki J, Imanishi E, Nagata S. Xkr8 phospholipid scrambling complex in apoptotic phosphatidylserine exposure. Proc Natl Acad Sci U S 2016;113: 9509–9514. doi:10.1073/pnas.1610403113

6. Segawa K, Kurata S, Yanagihashi Y, Brummelkamp TR, Matsuda F, Nagata S. Caspase-mediated cleavage of phospholipid flippase for apoptotic phosphatidylserine exposure. Science. 2014;344: 1164–1168. doi:10.1126/science.1252809

7. Nishi C, Toda S, Segawa K, Nagata S. Tim4-and MerTK-mediated engulfment of apoptotic cells by mouse resident peritoneal macrophages. Molecular and cellular biology. 2014 [cited 21 Mar 2014]. doi:10.1128/MCB.01394-13

8. Seitz HM, Camenisch TD, Lemke G, Earp HS, Matsushima GK. Macrophages and dendritic cells use different Axl/Mertk/Tyro3 receptors in clearance of apoptotic cells. Journal of immunology (Baltimore, Md: 1950). 2007;178: 5635–5642. doi:178/9/5635 [pii]

9. Sándor N, Lukácsi S, Ungai-Salánki R, Orgován N, Szabó B, Horváth R, et al. CD11c/CD18 Dominates Adhesion of Human Monocytes, Macrophages and Dendritic Cells over CD11b/CD18. Ng LG, editor. PLOS ONE. 2016;11: e0163120. doi:10.1371/journal.pone.0163120

10. Bain CC, Scott CL, Uronen-Hansson H, Gudjonsson S, Jansson O, Grip O, et al. Resident and pro-inflammatory macrophages in the colon represent alternative context-dependent fates of the same Ly6Chi monocyte precursors. Mucosal immunology. 2013;6: 498–510. doi:10.1038/mi.2012.89

11. Gardai SJ, McPhillips K a, Frasch SC, Janssen WJ, Starefeldt A, Murphy-Ullrich JE, et al. Cell-surface calreticulin initiates clearance of viable or apoptotic cells through trans-activation of LRP on the phagocyte. Cell. 2005;123: 321–34. doi:10.1016/j.cell.2005.08.032

12. Heit B, Kim H, Cosío G, Castaño D, Collins R, Lowell CA, et al. Multimolecular signaling complexes enable Syk-mediated signaling of CD36 internalization. Developmental cell. 2013;24: 372–83. doi:10.1016/j.devcel.2013.01.007

13. Leventis PA, Grinstein S. The distribution and function of phosphatidylserine in cellular membranes. Annu Rev Biophys. 2010;39: 407–427. doi:10.1146/annurev.biophys.093008.131234

14. Flannagan RS, Canton J, Furuya W, Glogauer M, Grinstein S. The phosphatidylserine receptor TIM4 utilizes integrins as coreceptors to effect phagocytosis. Molecular biology of the cell. 2014;25: 1511–22. doi:10.1091/mbc.E13-04-0212

15. Finnemann SC, Nandrot EF. MerTK activation during RPE phagocytosis in vivo requires alphaVbeta5 integrin. Advances in Experimental Medicine and Biology. 2006;572: 499–503. doi:10.1007/0-387-32442-9_69

16. Blackburn JWD, Lau DHC, Liu EY, Ellins J, Vrieze AM, Pawlak EN, et al. Soluble CD93 is an apoptotic cell opsonin recognized by αx β2. Eur J Immunol. 2019;49: 600–610. doi:10.1002/eji.201847801

17. Sadhu C, Ting HJ, Lipsky B, Hensley K, Garcia-Martinez LF, Simon SI, et al. CD11c/CD18: novel ligands and a role in delayed-type hypersensitivity. J Leukoc Biol. 2007;81: 1395–1403. doi:10.1189/jlb.1106680

18. Nagy-Baló Z, Kiss R, Menge A, Bödör C, Bajtay Z, Erdei A. Activated Human Memory B Lymphocytes Use CR4 (CD11c/CD18) for Adhesion, Migration, and Proliferation. Front Immunol. 2020;11: 565458. doi:10.3389/fimmu.2020.565458

19. Lukácsi S, Gerecsei T, Balázs K, Francz B, Szabó B, Erdei A, et al. The differential role of CR3 (CD11b/CD18) and CR4 (CD11c/CD18) in the adherence, migration and podosome formation of human macrophages and dendritic cells under inflammatory conditions. PLoS One. 2020;15: e0232432. doi:10.1371/journal.pone.0232432

20. McDowall A, Leitinger B, Stanley P, Bates PA, Randi AM, Hogg N. The I domain of integrin leukocyte function-associated antigen-1 is involved in a conformational change leading to high affinity binding to ligand intercellular adhesion molecule 1 (ICAM-1). J Biol Chem. 1998;273: 27396–27403. doi:10.1074/jbc.273.42.27396

21. Lefort CT, Hyun Y-M, Schultz JB, Law F-Y, Waugh RE, Knauf PA, et al. Outside-in signal transmission by conformational changes in integrin Mac-1. J Immunol. 2009;183: 6460–6468. doi:10.4049/jimmunol.0900983

22. Jaumouillé V, Cartagena-Rivera AX, Waterman CM. Coupling of β 2 integrins to actin by a mechanosensitive molecular clutch drives complement receptor-mediated phagocytosis. Nat Cell Biol. 2019;21: 1357–1369. doi:10.1038/s41556-019-0414-2

23. Freeman SA, Goyette J, Furuya W, Woods EC, Bertozzi CR, Bergmeier W, et al. Integrins Form an Expanding Diffusional Barrier that Coordinates Phagocytosis. Cell. 2016;164: 128–40. doi:10.1016/j.cell.2015.11.048

24. Choraghe RP, Kołodziej T, Buser A, Rajfur Z, Neumann AK. RHOA-mediated mechanical force generation through Dectin-1. J Cell Sci. 2020;133: jcs236166. doi:10.1242/jcs.236166

25. Bilsland CA, Diamond MS, Springer TA. The leukocyte integrin p150,95 (CD11c/CD18) as a receptor for iC3b. Activation by a heterologous beta subunit and localization of a ligand recognition site to the I domain. J Immunol. 1994;152: 4582–4589.

26. Jensen RK, Bajic G, Sen M, Springer TA, Vorup-Jensen T, Andersen GR. Complement Receptor 3 Forms a Compact High-Affinity Complex with iC3b. J Immunol. 2021;206: 3032–3042. doi:10.4049/jimmunol.2001208

27. Torres-Gomez A, Sanchez-Trincado JL, Toribio V, Torres-Ruiz R, Rodríguez-Perales S, Yáñez-Mó M, et al. RIAM-VASP Module Relays Integrin Complement Receptors in Outside-In Signaling Driving Particle Engulfment. Cells. 2020;9: E1166. doi:10.3390/cells9051166

28. DeBerge M, Yeap XY, Dehn S, Zhang S, Grigoryeva L, Misener S, et al. MerTK Cleavage on Resident Cardiac Macrophages Compromises Repair After Myocardial Ischemia Reperfusion Injury. Circ Res. 2017;121: 930–940. doi:10.1161/CIRCRESAHA.117.311327

29. Brea-Fernández AJ, Pomares E, Brión MJ, Marfany G, Blanco MJ, Sánchez-Salorio M, et al. Novel splice donor site mutation in MERTK gene associated with retinitis pigmentosa. The British journal of ophthalmology. 2008;92: 1419–1423. doi:10.1136/bjo.2008.139204

30. Binder MD, Fox AD, Merlo D, Johnson LJ, Giuffrida L, Calvert SE, et al. Common and Low Frequency Variants in MERTK Are Independently Associated with Multiple Sclerosis Susceptibility with Discordant Association Dependent upon HLA-DRB1*15:01 Status. PLoS genetics. 2016;12: e1005853. doi:10.1371/journal.pgen.1005853

31. Ma GZM, Stankovich J, Kilpatrick TJ, Binder MD, Field J. Polymorphisms in the receptor tyrosine kinase MERTK gene are associated with multiple sclerosis susceptibility. PloS one. 2011;6: e16964. doi:10.1371/journal.pone.0016964

32. Chen Y, Wang H, Qi N, Wu H, Xiong W, Ma J, et al. Functions of TAM RTKs in regulating spermatogenesis and male fertility in mice. Reproduction. 2009;138: 655–666. doi:10.1530/REP-09-0101

33. Shelby SJ, Colwill K, Dhe-Paganon S, Pawson T, Thompson D a. MERTK Interactions with SH2-Domain Proteins in the Retinal Pigment Epithelium. PloS one. 2013;8: e53964. doi:10.1371/journal.pone.0053964

34. Nandrot EF, Silva KE, Scelfo C, Finnemann SC. Retinal pigment epithelial cells use a MerTK-dependent mechanism to limit the phagocytic particle binding activity of αvβ5 integrin. Biol Cell. 2012;104: 326–341. doi:10.1111/boc.201100076

35. Wu Y, Singh S, Georgescu M-M, Birge RB. A role for Mer tyrosine kinase in alphavbeta5 integrin-mediated phagocytosis of apoptotic cells. Journal of Cell Science. 2005;118: 539–553. doi:10.1242/jcs.01632

36. Mao Y, Finnemann SC. Essential diurnal Rac1 activation during retinal phagocytosis requires αvβ5 integrin but not tyrosine kinases focal adhesion kinase or Mer tyrosine kinase. Mol Biol Cell. 2012;23: 1104– 1114. doi:10.1091/mbc.E11-10-0840

37. Dransfield I, Zagórska a, Lew ED, Michail K, Lemke G. Mer receptor tyrosine kinase mediates both tethering and phagocytosis of apoptotic cells. Cell Death and Disease. 2015;6: e1646. doi:10.1038/cddis.2015.18

38. Toda S, Segawa K, Nagata S. MerTK-mediated engulfment of pyrenocytes by central macrophages in erythroblastic islands. Blood. 2014;123: 3963–71. doi:10.1182/blood-2014-01-547976

39. Hu B, Jennings JH, Sonstein J, Floros J, Todt JC, Polak T, et al. Resident murine alveolar and peritoneal macrophages differ in adhesion of apoptotic thymocytes. Am J Respir Cell Mol Biol. 2004;30: 687– 693. doi:10.1165/rcmb.2003-0255OC

40. Zhang W, Zhang D, Stashko MA, DeRyckere D, Hunter D, Kireev D, et al. Pseudo-Cyclization through Intramolecular Hydrogen Bond Enables Discovery of Pyridine Substituted Pyrimidines as New Mer Kinase Inhibitors. J Med Chem. 2013;56: 9683– 9692. doi:10.1021/jm401387j

41. Evans AL, Blackburn JWD, Taruc K, Kipp A, Dirk BS, Hunt NR, et al. Antagonistic Coevolution of MER Tyrosine Kinase Expression and Function. Molecular biology and evolution. 2017;34: 1613–1628. doi:10.1093/molbev/msx102

42. Freeman SA, Grinstein S. Phagocytosis: receptors, signal integration, and the cytoskeleton. Immunological reviews. 2014;262: 193–215. doi:10.1111/imr.12212

43. Kovari DT, Wei W, Chang P, Toro J-S, Beach RF, Chambers D, et al. Frustrated Phagocytic Spreading of J774A-1 Macrophages Ends in Myosin II-Dependent Contraction. Biophysical journal. 2016;111: 2698–2710. doi:10.1016/j.bpj.2016.11.009

44. Lin J, Kurilova S, Scott BL, Bosworth E, Iverson BE, Bailey EM, et al. TIRF imaging of Fc gamma receptor microclusters dynamics and signaling on macrophages during frustrated phagocytosis. BMC immunology. 2016;17: 5. doi:10.1186/s12865-016-0143-2

45. Crowley MT, Costello PS, Fitzer-Attas CJ, Turner M, Meng F, Lowell C, et al. A critical role for Syk in signal transduction and phagocytosis mediated by Fcgamma receptors on macrophages. J Exp Med. 1997;186: 1027–1039.

46. Jaqaman K, Kuwata H, Touret N, Collins R, Trimble WS, Danuser G, et al. Cytoskeletal Control of CD36 Diffusion Promotes Its Receptor and Signaling Function. Cell. 2011;146: 593–606. doi:10.1016/j.cell.2011.06.049

47. Kusumi A, Nakada C, Ritchie K, Murase K, Suzuki K, Murakoshi H, et al. Paradigm shift of the plasma membrane concept from the two-dimensional continuum fluid to the partitioned fluid: high-speed single-molecule tracking of membrane molecules. Annual review of biophysics and biomolecular structure. 2005;34: 351–78. doi:10.1146/annurev.biophys.34.040204.144637

48. Klapholz B, Brown NH. Talin-the master of integrin adhesions. Journal of cell science. 2017;130: 2435–2446. doi:10.1242/jcs.190991

49. Freeman SA, Vega A, Riedl M, Collins RF, Ostrowski PP, Woods EC, et al. Transmembrane Pickets Connect Cyto-and Pericellular Skeletons Forming Barriers to Receptor Engagement. Cell. 2018;172: 305–312.e10. doi:10.1016/j.cell.2017.12.023

50. Rose M, Hirmiz N, Moran-Mirabal JM, Fradin C. Lipid Diffusion in Supported Lipid Bilayers: A Comparison between Line-Scanning Fluorescence Correlation Spectroscopy and Single-Particle Tracking. Membranes (Basel). 2015;5: 702–721. doi:10.3390/membranes5040702

51. Marchetti L, Callegari A, Luin S, Signore G, Viegi A, Beltram F, et al. Ligand signature in the membrane dynamics of single TrkA receptor molecules. J Cell Sci. 2013;126: 4445–4456. doi:10.1242/jcs.129916

52. Su Z, Dhusia K, Wu Y. A multiscale study on the mechanisms of spatial organization in ligand-receptor interactions on cell surfaces. Comput Struct Biotechnol J. 2021;19: 1620–1634. doi:10.1016/j.csbj.2021.03.024

53. Feng W, Yasumura D, Matthes MT, LaVail MM, Vollrath D. Mertk triggers uptake of photoreceptor outer segments during phagocytosis by cultured retinal pigment epithelial cells. The Journal of biological chemistry. 2002;277: 17016–22. doi:10.1074/jbc.M107876200

54. Cai B, Thorp EB, Doran AC, Sansbury BE, Daemen MJAP, Dorweiler B, et al. MerTK receptor cleavage promotes plaque necrosis and defective resolution in atherosclerosis. The Journal of clinical investigation. 2017;127: 1–5. doi:10.1172/JCI90520

55. Thorp E, Cui D, Schrijvers DM, Kuriakose G, Tabas I. Mertk receptor mutation reduces efferocytosis efficiency and promotes apoptotic cell accumulation and plaque necrosis in atherosclerotic lesions of apoe-/-mice. Arteriosclerosis, thrombosis, and vascular biology. 2008;28: 1421–8. doi:10.1161/ATVBAHA.108.167197

56. Hurtado BBB, Abasolo N, Muñoz X, García N, Benavente Y, Rubio F, et al. Association study between polymorphims in GAS6-TAM genes and carotid atherosclerosis. Thrombosis and Haemostasis. 2010;104: 592–8. doi:10.1160/TH09-11-0787

57. Vorselen D, Barger SR, Wang Y, Cai W, Theriot JA, Gauthier NC, et al. Phagocytic “teeth” and myosin-II “jaw” power target constriction during phagocytosis. Elife. 2021;10: e68627. doi:10.7554/eLife.68627

58. Barger SR, Vorselen D, Gauthier NC, Theriot JA, Krendel M. F-actin organization and target constriction during primary macrophage phagocytosis is balanced by competing activity of myosin-I and myosin-II. MBoC. 2022;33: br24. doi:10.1091/mbc.E22-06-0210

59. Francis EA, Xiao H, Teng LH, Heinrich V. Mechanisms of frustrated phagocytic spreading of human neutrophils on antibody-coated surfaces. Biophysical Journal. 2022;121: 4714–4728. doi:10.1016/j.bpj.2022.10.016

60. Cordoba S-P, Choudhuri K, Zhang H, Bridge M, Basat AB, Dustin ML, et al. The large ectodomains of CD45 and CD148 regulate their segregation from and inhibition of ligated T-cell receptor. Blood. 2013;121: 4295–4302. doi:10.1182/blood-2012-07-442251

61. Thiagarajan P, Tait JF. Binding of annexin V/placental anticoagulant protein I to platelets. Evidence for phosphatidylserine exposure in the procoagulant response of activated platelets. J Biol Chem. 1990;265: 17420–3. doi:2145274

62. Lentz BR. Exposure of platelet membrane phosphatidylserine regulates blood coagulation. Progress in Lipid Research. 2003;42: 423–438. doi:10.1016/S0163-7827(03)00025-0

63. Smrz D, Dráberová L, Dráber P. Non-apoptotic phosphatidylserine externalization induced by engagement of glycosylphosphatidylinositol-anchored proteins. J Biol Chem. 2007;282: 10487–10497. doi:10.1074/jbc.M611090200

64. Elliott JI, Surprenant A, Marelli-Berg FM, Cooper JC, Cassady-Cain RL, Wooding C, et al. Membrane phosphatidylserine distribution as a non-apoptotic signalling mechanism in lymphocytes. Nat Cell Biol. 2005;7: 808–816. doi:10.1038/ncb1279

65. van den Eijnde SM, van den Hoff MJ, Reutelingsperger CP, van Heerde WL, Henfling ME, Vermeij-Keers C, et al. Transient expression of phosphatidylserine at cell-cell contact areas is required for myotube formation. J Cell Sci. 2001;114: 3631– 3642. doi:10.1242/jcs.114.20.3631

66. Dillon SR, Mancini M, Rosen A, Schlissel MS. Annexin V binds to viable B cells and colocalizes with a marker of lipid rafts upon B cell receptor activation. J Immunol. 2000;164: 1322–1332. doi:10.4049/jimmunol.164.3.1322

67. Audo R, Hua C, Hahne M, Combe B, Morel J, Daien CI. Phosphatidylserine Outer Layer Translocation Is Implicated in IL-10 Secretion by Human Regulatory B Cells. PLoS One. 2017;12: e0169755. doi:10.1371/journal.pone.0169755

68. Fischer K, Voelkl S, Berger J, Andreesen R, Pomorski T, Mackensen A. Antigen recognition induces phosphatidylserine exposure on the cell surface of human CD8+ T cells. Blood. 2006;108: 4094–4101. doi:10.1182/blood-2006-03-011742

69. Stowell SR, Karmakar S, Stowell CJ, Dias-Baruffi M, McEver RP, Cummings RD. Human galectin-1, −2, and −4 induce surface exposure of phosphatidylserine in activated human neutrophils but not in activated T cells. Blood. 2007;109: 219–227. doi:10.1182/blood-2006-03-007153

70. Vernon-Wilson EF, Kee WJ, Willis AC, Barclay AN, Simmons DL, Brown MH. CD47 is a ligand for rat macrophage membrane signal regulatory protein SIRP (OX41) and human SIRPalpha 1. Eur J Immunol. 2000;30: 2130–2137. doi:10.1002/1521-4141(2000)30:8<2130::AID-IMMU2130>3.0.CO;2-8

71. Morrissey MA, Kern N, Vale RD. CD47 Ligation Repositions the Inhibitory Receptor SIRPA to Suppress Integrin Activation and Phagocytosis. Immunity. 2020;53: 290–302.e6. doi:10.1016/j.immuni.2020.07.008

72. Lv Z, Bian Z, Shi L, Niu S, Ha B, Tremblay A, et al. Loss of Cell Surface CD47 Clustering Formation and Binding Avidity to SIRPα Facilitate Apoptotic Cell Clearance by Macrophages. J Immunol. 2015;195: 661–671. doi:10.4049/jimmunol.1401719

73. Hirano Y, Yang W-L, Aziz M, Zhang F, Sherry B, Wang P. MFG-E8-derived peptide attenuates adhesion and migration of immune cells to endothelial cells. Journal of Leukocyte Biology. 2017;101: 1201– 1209. doi:10.1189/jlb.3A0416-184RR

74. Chigaev A, Blenc AM, Braaten JV, Kumaraswamy N, Kepley CL, Andrews RP, et al. Real Time Analysis of the Affinity Regulation of α4-Integrin: THE PHYSIOLOGICALLY ACTIVATED RECEPTOR IS INTERMEDIATE IN AFFINITY BETWEEN RESTING AND Mn2+ OR ANTIBODY ACTIVATION*. Journal of Biological Chemistry. 2001;276: 48670–48678. doi:10.1074/jbc.M103194200

75. Li J, Yan J, Springer TA. Low-affinity integrin states have faster ligand-binding kinetics than the high-affinity state. Fässler R, Faraldo-Gómez JD, Fässler R, editors. eLife. 2021;10: e73359. doi:10.7554/eLife.73359

76. Connors WL, Jokinen J, White DJ, Puranen JS, Kankaanpää P, Upla P, et al. Two synergistic activation mechanisms of alpha2beta1 integrin-mediated collagen binding. J Biol Chem. 2007;282: 14675– 14683. doi:10.1074/jbc.M700759200

77. Kiefer F, Brumell J, Al-Alawi N, Latour S, Cheng a, Veillette a, et al. The Syk protein tyrosine kinase is essential for Fcgamma receptor signaling in macrophages and neutrophils. Molecular and cellular biology. 1998;18: 4209–20.

78. Walbaum S, Ambrosy B, Schütz P, Bachg AC, Horsthemke M, Leusen JHW, et al. Complement receptor 3 mediates both sinking phagocytosis and phagocytic cup formation via distinct mechanisms. Journal of Biological Chemistry. 2021;296: 100256. doi:10.1016/j.jbc.2021.100256

79. Shiratsuchi H, Basson MD. Extracellular pressure stimulates macrophage phagocytosis by inhibiting a pathway involving FAK and ERK. American Journal of Physiology-Cell Physiology. 2004;286: C1358–C1366. doi:10.1152/ajpcell.00553.2003

80. Sayedyahossein S, Nini L, Irvine TS, Dagnino L. Essential role of integrin-linked kinase in regulation of phagocytosis in keratinocytes. FASEB J. 2012;26: 4218–4229. doi:10.1096/fj.12-207852

81. Antonieta Cote-Vélez MJ, Ortega E, Ortega A. Involvement of pp125FAK and p60SRC in the signaling through Fc gamma RII-Fc gamma RIII in murine macrophages. Immunol Lett. 2001;78: 189–194. doi:10.1016/s0165-2478(01)00251-6

82. Toda S, Hanayama R, Nagata S. Two-step engulfment of apoptotic cells. Molecular and cellular biology. 2012;32: 118–25. doi:10.1128/MCB.05993-11

83. Moon B, Lee J, Lee S-A, Min C, Moon H, Kim D, et al. Mertk Interacts with Tim-4 to Enhance Tim-4-Mediated Efferocytosis. Cells. 2020;9: E1625. doi:10.3390/cells9071625

84. Brouillette RB, Phillips EK, Patel R, Mahauad-Fernandez W, Moller-Tank S, Rogers KJ, et al. TIM-1 Mediates Dystroglycan-Independent Entry of Lassa Virus. J Virol. 2018;92: e00093–18. doi:10.1128/JVI.00093-18

85. Penberthy KK, Rival C, Shankman LS, Raymond MH, Zhang J, Perry JSA, et al. Context-dependent compensation among phosphatidylserine-recognition receptors. Sci Rep. 2017;7: 14623. doi:10.1038/s41598-017-15191-1

86. Park D, Tosello-Trampont A-C, Elliott MR, Lu M, Haney LB, Ma Z, et al. BAI1 is an engulfment receptor for apoptotic cells upstream of the ELMO/Dock180/Rac module. Nature. 2007;450: 430–4. doi:10.1038/nature06329

87. Lee WL, Cosio G, Ireton K, Grinstein S. Role of CrkII in Fcgamma receptor-mediated phagocytosis. J Biol Chem. 2007;282: 11135–11143. doi:10.1074/jbc.M700823200

88. Jiang Y, Zhang Y, Leung JY, Fan C, Popov KI, Su S, et al. MERTK mediated novel site Akt phosphorylation alleviates SAV1 suppression. Nat Commun. 2019;10: 1515. doi:10.1038/s41467-019-09233-7

89. Georgescu MM, Kirsch KH, Shishido T, Zong C, Hanafusa H. Biological effects of c-Mer receptor tyrosine kinase in hematopoietic cells depend on the Grb2 binding site in the receptor and activation of NF-kappaB. Molecular and cellular biology. 1999;19: 1171– 81.

90. Garçon F, Okkenhaug K. PI3Kδ promotes CD4+ T-cell interactions with antigen-presenting cells by increasing LFA-1 binding to ICAM-1. Immunol Cell Biol. 2016;94: 486–495. doi:10.1038/icb.2016.1

91. Vachon E, Martin R, Kwok V, Cherepanov V, Chow C-WC-WC-W, Doerschuk CM, et al. CD44-mediated phagocytosis induces inside-out activation of complement receptor-3 in murine macrophages. Blood. 2007;110: 4492–502. doi:10.1182/blood-2007-02-076539

92. Schindelin J, Arganda-Carreras I, Frise E, Kaynig V, Longair M, Pietzsch T, et al. Fiji: an open-source platform for biological-image analysis. Nature methods. 2012;9: 676–82. doi:10.1038/nmeth.2019

93. Yin C, Vrieze AM, Rosoga M, Akingbasote J, Pawlak EN, Jacob RA, et al. Efferocytic Defects in Early Atherosclerosis Are Driven by GATA2 Overexpression in Macrophages. Front Immunol. 2020;11: 594136. doi:10.3389/fimmu.2020.594136

94. Pereira PM, Albrecht D, Culley S, Jacobs C, Marsh M, Mercer J, et al. Fix Your Membrane Receptor Imaging: Actin Cytoskeleton and CD4 Membrane Organization Disruption by Chemical Fixation. Front Immunol. 2019;10: 675. doi:10.3389/fimmu.2019.00675

95. Caetano Crowley FA, Heit B, Ferguson SSG. Super-Resolution Imaging of G Protein-Coupled Receptors Using Ground State Depletion Microscopy. Methods Mol Biol. 2019;1947: 323–336. doi:10.1007/978-1-4939-9121-1_18

96. Caetano FA, Dirk BS, Tam JHK, Cavanagh PC, Goiko M, Ferguson SSG, et al. MIiSR: Molecular Interactions in Super-Resolution Imaging Enables the Analysis of Protein Interactions, Dynamics and Formation of Multi-protein Structures. Mac Gabhann F, editor. PLoS computational biology. 2015;11: e1004634. doi:10.1371/journal.pcbi.1004634

97. Taruc K, Yin C, Wootton DG, Heit B. Quantification of Efferocytosis by Single-cell Fluorescence Microscopy. Journal of visualized experiments: JoVE. 2018. doi:10.3791/58149

98. Heit B, Robbins SM, Downey CM, Guan Z, Colarusso P, Miller BJ, et al. PTEN functions to “prioritize” chemotactic cues and prevent “distraction” in migrating neutrophils. Nature immunology. 2008;9: 743–52. doi:10.1038/ni.1623

99. Heit B, Tavener S, Raharjo E, Kubes P. An intracellular signaling hierarchy determines direction of migration in opposing chemotactic gradients. The Journal of cell biology. 2002;159: 91–102. doi:10.1083/jcb.200202114

100. Yin C, Kim Y, Argintaru D, Heit B. Rab17 mediates differential antigen sorting following efferocytosis and phagocytosis. Cell death & disease. 2016;7: e2529. doi:10.1038/cddis.2016.431

101. van Rheenen J, Langeslag M, Jalink K. Correcting Confocal Acquisition to Optimize Imaging of Fluorescence Resonance Energy Transfer by Sensitized Emission. Biophysical Journal. 2004;86: 2517–2529. doi:10.1016/S0006-3495(04)74307-6

102. Tepperman A, Zheng DJ, Taka MA, Vrieze A, Le Lam A, Heit B. Customizable live-cell imaging chambers for multimodal and multiplex fluorescence microscopy. Biochemistry and cell biology = Biochimie et biologie cellulaire. 2020;98: 612–623. doi:10.1139/bcb-2020-0064

103. Bolte S, Cordelières FP. A guided tour into subcellular colocalization analysis in light microscopy. Journal of microscopy. 2006;224: 213–32. doi:10.1111/j.1365-2818.2006.01706.x

104. Armstrong SM, Sugiyama MG, Fung KYY, Gao Y, Wang C, Levy AS, et al. A novel assay uncovers an unexpected role for SR-BI in LDL transcytosis. Cardiovascular Research. 2015;108: 268–277. doi:10.1093/cvr/cvv218

105. Goiko M, de Bruyn JR, Heit B. Short-Lived Cages Restrict Protein Diffusion in the Plasma Membrane. Scientific reports. 2016;6: 34987. doi:10.1038/srep34987

106. Goiko M, de Bruyn JR, Heit B. Membrane Diffusion Occurs by Continuous-Time Random Walk Sustained by Vesicular Trafficking. Biophysical journal. 2018;114: 2887–2899. doi:10.1016/j.bpj.2018.04.024

107. Jaqaman K, Loerke D, Mettlen M, Kuwata H, Grinstein S, Schmid SL, et al. Robust single-particle tracking in live-cell time-lapse sequences. Nature methods. 2008;5: 695–702. doi:10.1038/nmeth.1237

108. Ferrari R, Manfroi AJ, Young WR. Strongly and weakly self-similar diffusion. Physica D: Nonlinear Phenomena. 2001;154: 111–137. doi:10.1016/S0167-2789(01)00234-2

