## Supplemental Figures 1-5 for "MERTK Coordinates Efferocytosis by Regulating Integrin Localization and Activation"

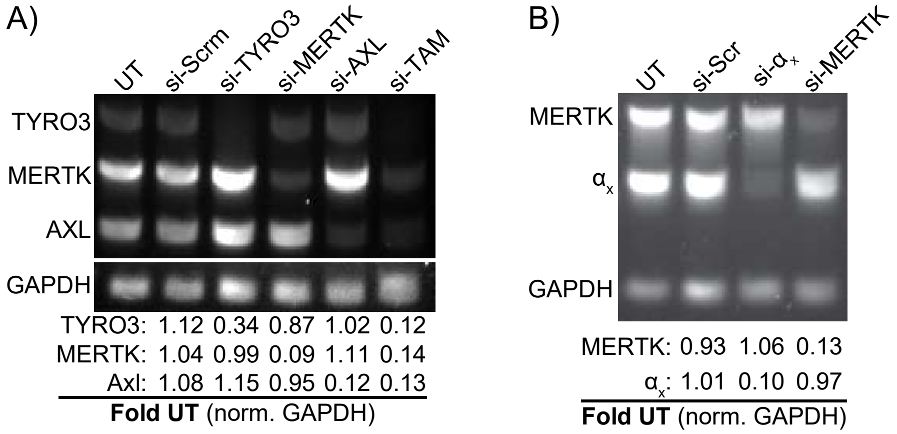


**Supplemental Figure 1.** Confirmation of siRNA depletion by semi-quantitative RT-PCR. **A)** Depletion of the TAM receptors individually (si-MERTK, si-Axl and si-Tyro3) and combined (si-TAM) compared to non-transfected (UT) and cells treated with a scrambled (non-targeting) siRNA (si-Scrm). **B)** siRNA knockdown of MERTK and $\alpha_x$ integrin. Data is representative of 3 independent experiments.

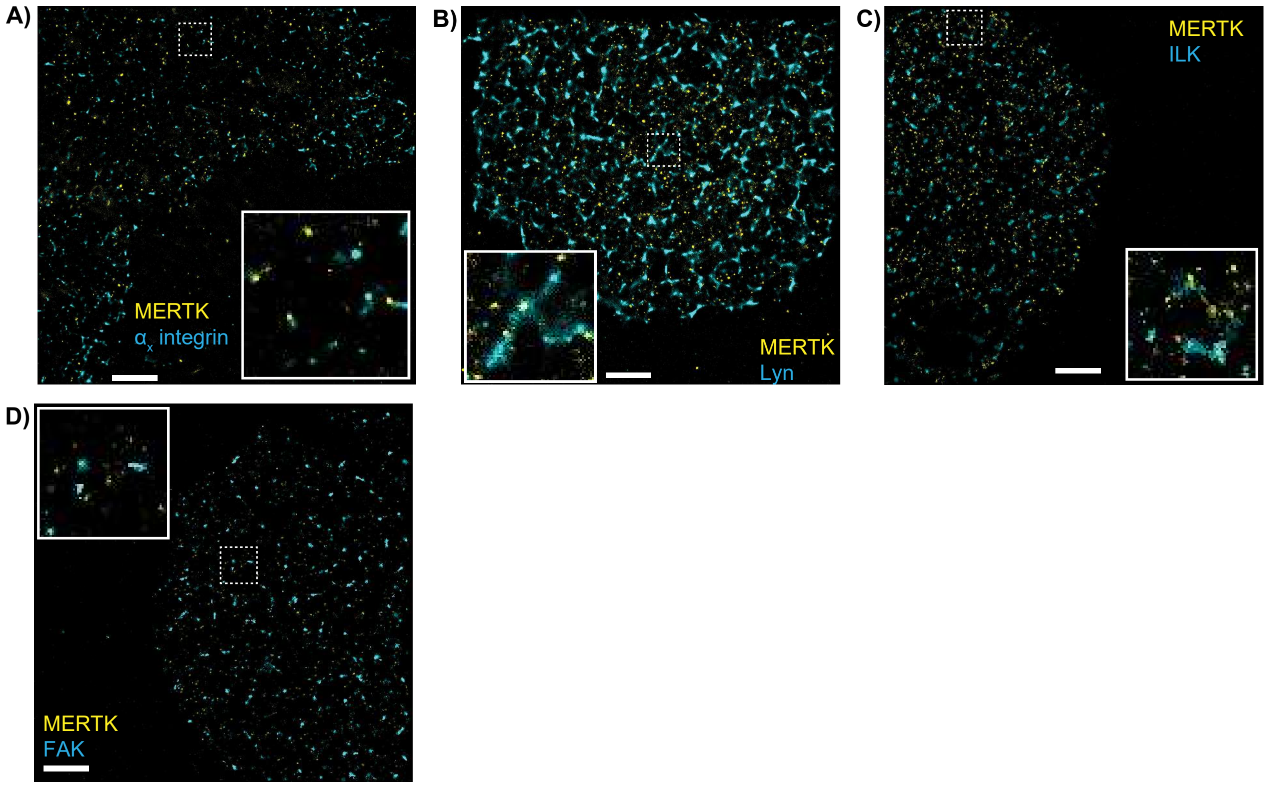


**Supplemental Figure 2.** Ground state depletion microscopy images showing the distribution on the surface of macrophages of MERTK (yellow) and $\alpha_x$ integrin (A), Lyn (B), ILK (C), and FAK (D). Inserts show regions within the dotted boxes, scale bars are 2 μm. Data is representative of 3-5 independent experiments.

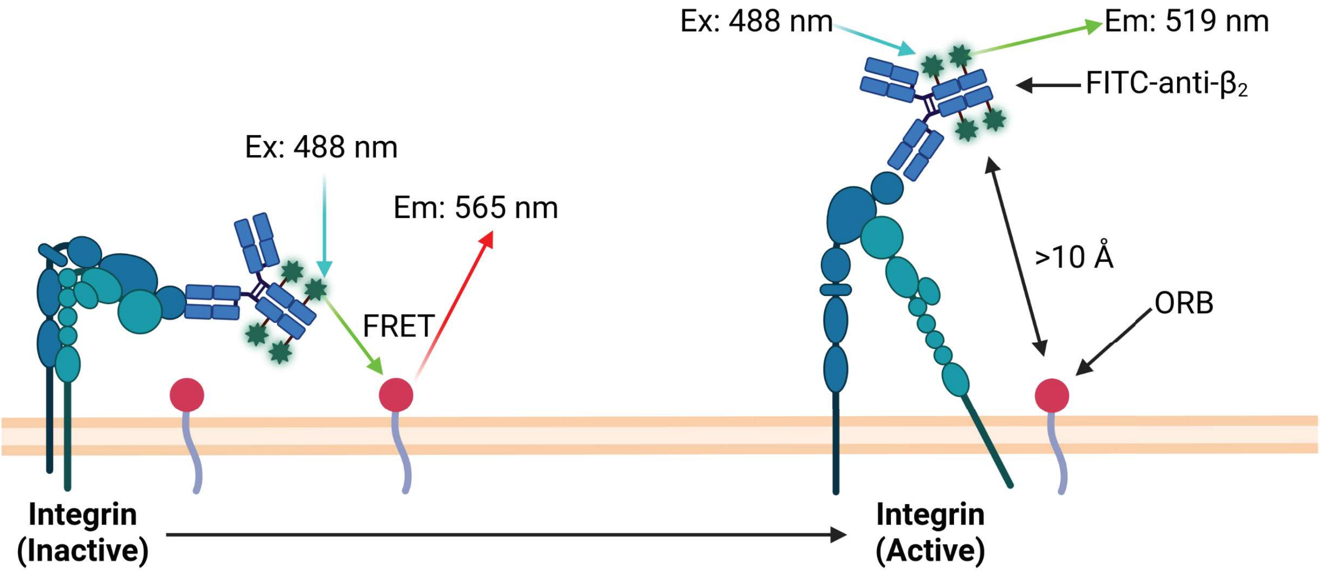


**Supplemental Figure 3.** Model of FRET-based integrin activation assay. When the β2 integrin is in the inactive (bent) conformation, the FITC-labeled anti-$\beta_2$ headgroup-binding antibody TS1/18 will be positioned near the plasma membrane, where the excitation energy of FITC can be transferred to octadecylrhodamine B (ORB) embedded in the macrophage membrane. This results in the presence of ORB emission (565 nm) when FITC is excited using 488 nm excitation. Activation of the integrin causes a conformational change which will move the headgroup away from the plasma membrane, thereby positioning the TS1/18 antibody beyond the 10Å limit of FRET energy transfer. This causes a loss of the ORB fluorescent signal, and a concordant increase in the FITC emission at 519 nm.

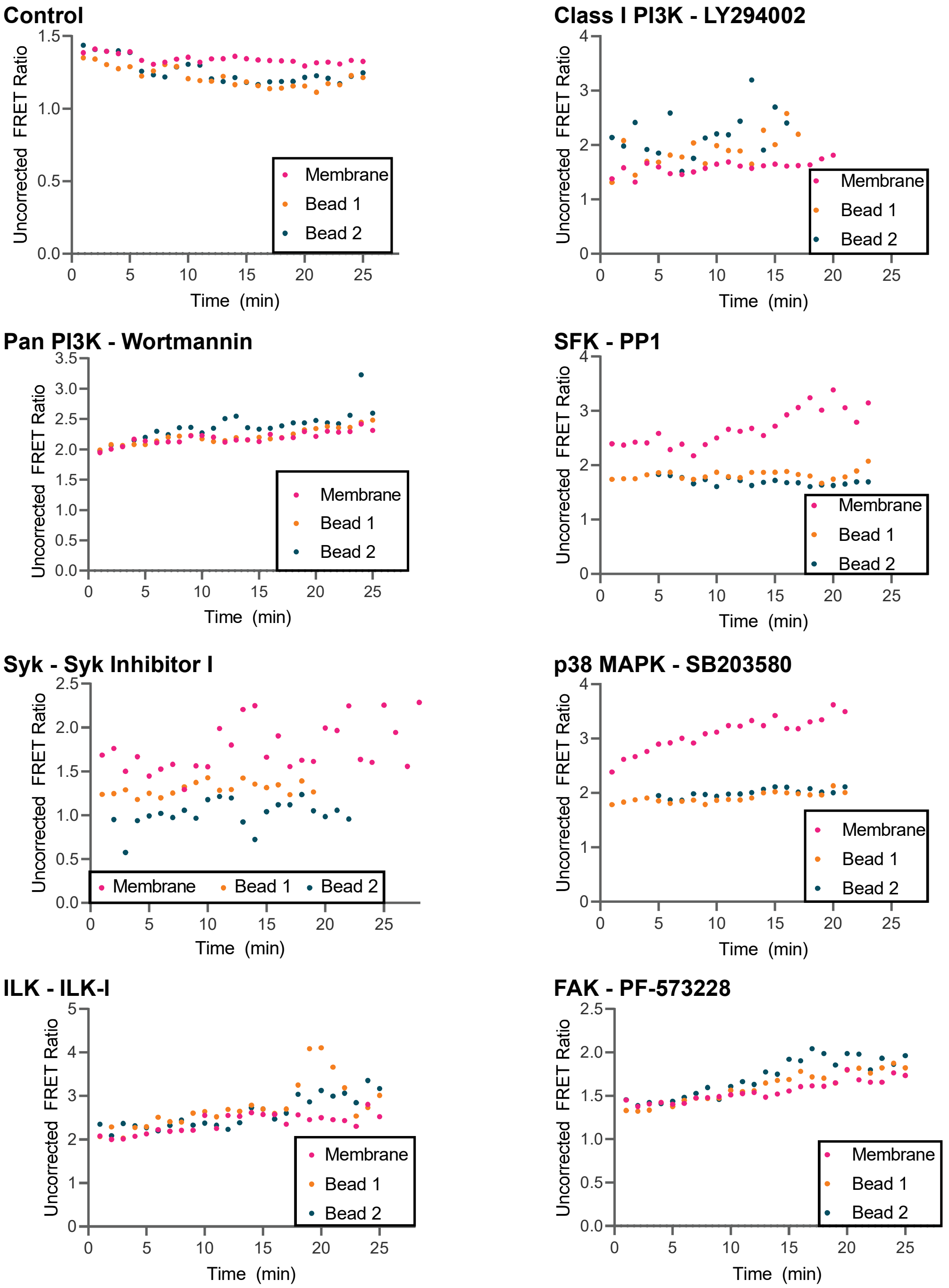


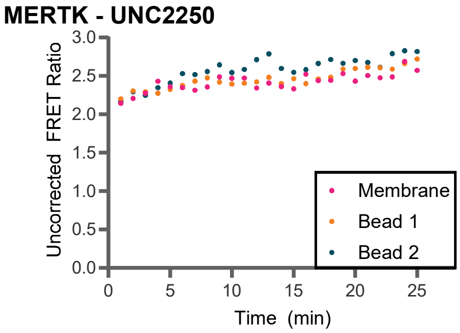


**Supplemental Figure 4**. $\beta_2$ integrin activation dynamics, comparing the uncorrected FRET ratio between FITC and ORB of the bulk membrane (Membrane) versus the membrane in-contact with two representative Gas6-opsonized apoptotic cell mimics (Bead 1/Bead 2). $\beta_2$ integrin activation within the phagocytic cup is apparent as a decrease in the FRET ratio of the membrane contacting the beads relative to the bulk membrane. Cells are pretreated with the indicated inhibitors or a vehicle control. Data is representative of a minimum of 30 cells imaged in three separate experiments.

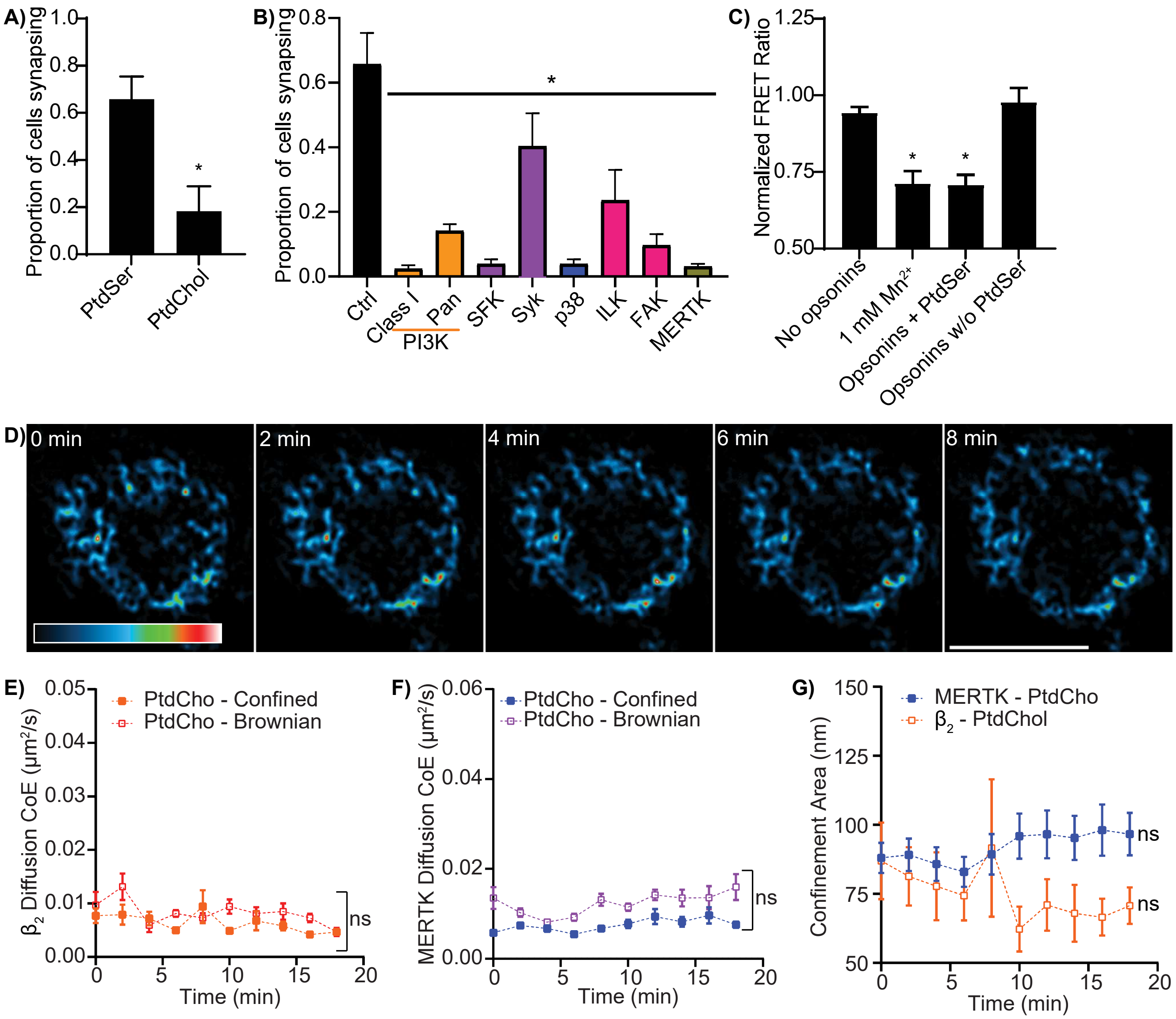


**Supplemental Figure 5. A)** Portion of macrophages forming a synapse on Gas6 and MFG-E8 opsonized planar lipid bilayers containing either 20:80 PtdSer:PtdChol (PtdSer) versus 100% PtdChol. **B)** Effect of PI3K (LY & Wort), SFK (PP1 & SykI), p38 MAPK (SB), ILK (ILK-I), FAK (PF) and MERTK (UNC) inhibition, or a veihicle control (Ctrl) on the portion of THP1 macrophages which form a synapse when in contact with a planar apoptotic cell mimic. **C)** Activation of $\beta_2$ integrins, as quantified by FRET microscopy, on planar apoptotic mimics containing 20:80 PtdSer:PtdChol lacking opsonins (no opsonins), lacking opsonins and stimulated with manganese, opsonized with Gas6 and MFG-E8, or on a mimic containing 100% PtdChol that is opsonized with Gas6 and MFG-E8 (Opsonins w/o PtdSer). **D)** Actin distribution on a macrophage interacting with a 100% PtdChol containing lipid bilayer. **E-F)** Diffusion coefficient of (E) $\beta_2$ integrin and (F) MERTK on a 100% PtdChol containing lipid bilayer. **G)** Confinement area size of MERTK and $\beta_2$ integrin on macrophages interacting with a 100% PtdChol containing lipid bilayer. Data are plotted as mean ± SEM, n = 3 to 5. * $p < 0.05$ compared to PtdSer (A), UT (B), no opsonins (C), or T=0 (E-G), Students T-test (A) or Kruskal-Wallis test with Dunn's multiple comparisons test.
